# Deconstructing the Mapper algorithm to extract richer topological and temporal features from functional neuroimaging data

**DOI:** 10.1101/2023.10.13.562304

**Authors:** Daniel Haşegan, Caleb Geniesse, Samir Chowdhury, Manish Saggar

## Abstract

Capturing and tracking large-scale brain activity dynamics holds the potential to deepen our understanding of cognition. Previously, tools from Topological Data Analysis, especially Mapper, have been successfully used to mine brain activity dynamics at the highest spatiotemporal resolutions. Even though it is a relatively established tool within the field of Topological Data Analysis, Mapper results are highly impacted by parameter selection. Given that non-invasive human neuroimaging data (e.g., from fMRI) is typically fraught with artifacts and no gold standards exist regarding “true” state transitions, we argue for a thorough examination of Mapper parameter choices to better reveal their impact. Using synthetic data (with known transition structure) and real fMRI data, we explore a variety of parameter choices for each Mapper step, thereby providing guidance and heuristics for the field. We also release our parameter-exploration toolbox as a software package to make it easier for scientists to investigate and apply Mapper on any dataset.

## 1. Introduction

A main interest in neuroscience research is understanding the relationship between brain dynamics and behavior. Due to the high dimensionality and complexity of recorded neuronal data, computational methods have been developed to capture and track brain dynamics. While there are many available methods to quantify brain dynamics (Chang and Glover 2010; Liu and Duyn 2013; Xu and Lindquist 2015; Shine et al. 2016), with a few exceptions, most of them require collapsing (or selecting) data in space, time or across people at the outset (Saggar et al. 2018, 2022). To capture specific individual transitions in brain activity at the highest spatiotemporal resolutions without necessarily averaging (or selecting) data at the outset, the Topological Data Analysis based Mapper approach was developed (Singh et al. 2007; Saggar et al. 2018). The Mapper approach is typically used to characterize the “shape” of the underlying dataset as a graph (a.k.a. shape graph). Further, a priori knowledge about the number of whole-brain configurations is unnecessary, and Mapper does not impose strict assumptions about the mutual exclusivity of brain states (Baker et al. 2014).

Previously, Mapper has been applied to capture transitions in task-evoked (Saggar et al. 2018; Geniesse et al. 2019; Geniesse, Chowdhury, and Saggar 2022; M. Zhang, Chowdhury, and Saggar 2022) as well as intrinsic brain activity (Saggar et al. 2022). Mapper was also used to examine changes in brain dynamics associated with pharmacological interventions (Saggar et al. 2022). Even in domains beyond neuroimaging, Mapper has also been successfully utilized (Skaf and Laubenbacher 2022; Lum et al. 2013; Yao et al. 2009; Nicolau, Levine, and Carlsson 2011). While Mapper has been applied to neuroimaging data in the past, the parameter choices for Mapper have not been fully explored. Theoretical work has proposed a data-driven selection of mapper parameters (Carriere, Michel, and Oudot 2018; Chalapathi, Zhou, and Wang 2021), but the algorithms are limited to 1-dimensional covers, requiring more work for extending it to neuroimaging datasets that need higher dimensional covers. Current approaches to parameter selection on neuroimaging data are based on heuristics and educated guesses (Geniesse, Chowdhury, and Saggar 2022). To better understand the effect of parameter selection on neuroimaging data, we systematically deconstruct each Mapper step to reveal the impact of different parameter choices at each step. We also provide software tools for performing similar parameter explorations to facilitate wider applications of Mapper.

In a typical application of Mapper to study neural dynamics, after standard preprocessing steps, the high-dimensional data is fed to the pipeline as a 2D matrix, where rows correspond to individual time frames and columns correspond to regional activations. The Mapper pipeline consists of five main steps (**Fig 1**). First, a distance metric is picked to define the relationship between each row element in the original high-dimensional space. Second, the filter function embeds the data into a lower dimension. Third, overlapping low-dimensional binning is performed to allow for compression, putatively increasing reliability (by reducing noise-related perturbations). Fourth, partial clustering within each bin is performed, where the original distances between data points are used for coalescing (or separating) those points into graph nodes, allowing for the partial recovery of the information loss incurred due to the filter function (the dimensionality reduction). Lastly, nodes from different bins are connected if any data points are shared between them to generate a graphical representation of the data landscape. As a result, the topological information of the input data is represented as a “shape graph,” denoting the dynamical trajectory through recurring states.

**Figure 1:**
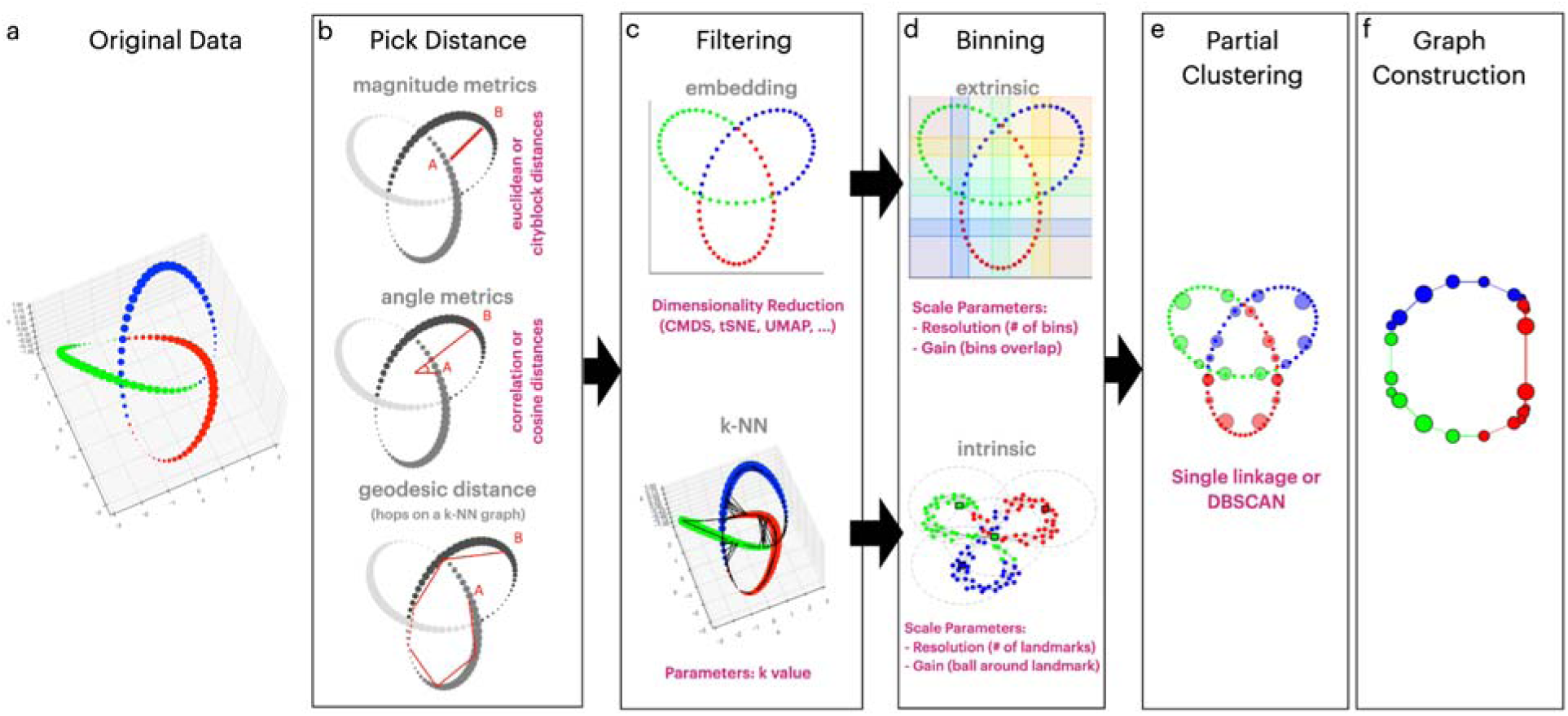
Mapper steps on synthetic Trefoil Knot. **(a)** The trefoil knot dataset contains sampled 3-dimensional points that are represented as dots. The true shape of this data is a closed loop. The points are colored to track their transformation in subsequent Mapper algorithm steps. **(b)** The first step of the Mapper algorithm is selecting a distance metric and optionally computing pairwise distances between all data points. One chooses between a magnitude metric such as Euclidean or Cityblock (Manhattan) distances, an angle metric such as Cosine or Correlation distance, or a geodesic metric based on a constructed k-Nearest Neighbor (k-NN) graph with an associated distance metric. The red lines between points A and B signify a schematic representation of the metric choice. **(c)** As a second step, the pairwise similarity matrix is projected to a reduced space (using a filter function) either through a dimensionality reduction algorithm or by selecting the k-Nearest Neighbors (k-NN) graph. When using a dimensionality reduction technique such as Classical Multidimensional Scaling (CMDS) or t-distributed stochastic neighbor embedding (t-SNE), the algorithm represents the sampled points in a lower dimensional (2 dimensions) embedding. Alternatively, using a k-NN algorithm, each point contains a list of neighbors as represented by black lines in the figure, connecting adjacent and distanced points based on the k-value used. The lens (or filter) choice determines the binning strategy, as the black arrows between the (c) and (d) boxes indicate. **(d)** The binning step segments the reduced space of points into coherent regions that create a cover of the lens (filter). An embedding filter requires the extrinsic binning choice, where the space of points is split into overlapping bins (in the figure, we use 36 rectangular bins, denoting a resolution of 6 bins per dimension; and overlap of bins of 20%, denoted as gain). For a k-NN filter, the intrinsic binning step selects points as landmarks and segments the space as distances from the picked landmarks (in the figure, we used four landmarks, denoting a resolution of 4, with the gain as the distance from a landmark). **(e)** As the partial clustering step of the Mapper algorithm, the points in each bin are clustered into groups using the single linkage clustering algorithm or Density-based spatial clustering of applications with noise (DBSCAN). Partial (within bin) clustering is done in the original high-dimensional space to reduce the information lost due to embedding. The clustered groups will represent the nodes in the constructed graph. **(f)** As the final step of the algorithm, the nodes are linked by edges, created based on shared data points between the clusters, creating the Mapper “shape graph.”

Although Mapper has successfully revealed brain dynamics at rest and task-evoked states, the algorithm’s parameter choices and their impact on the final resulting shape graphs are rarely scanned systematically. In this paper, using synthetic and real fMRI datasets, we examine multiple parameter choices for each deconstructed step of the algorithm to understand its final contribution to the shape graph of neural dynamics. We also explore the effect of noise on Mapper graphs and examine their resilience under high noise levels.

## 2. Methods

### 2.1 Mapper algorithm

The Mapper algorithm creates a graph representation that preserves the topological features of the inputted high-dimensional data (Lum et al. 2013; Singh et al. 2007). The input data is a set of measurements of N points with M features represented by a 2-dimensional NxM matrix. In **Fig. 1**, we outline the Mapper steps and results on a synthetic Trefoil knot dataset, where we sampled points with three features, the x, y, and z coordinates (**Fig. 1a**). For typical neuroimaging data, the time-by-regions matrix has data points collected at time intervals (repetition time or sampling rate) at specific anatomical locations (brain voxels or parcels, or scalp location).

We divided the Mapper algorithm into five consecutive steps: (i) pick a distance metric and optionally compute pairwise distances between all points (**Fig. 1b**); (ii) project the data into a reduced low-dimensional space (or create a k-NN graph in case of intrinsic binning later) (**Fig. 1c**); (iii) separate the space into overlapping bins (**Fig. 1d**); (iv) cluster points within bins in the low-dimensional space using information from the high-dimensional data, coalescing into nodes (**Fig. 1e**); and (v) link the nodes across bins if they share any datapoints (**Fig. 1f**). The result is a “shape” graph where each node represents one or more rows (or time points), and an edge represents shared rows between nodes.

While many parameter choices will extract the main topological features of the input data, some combinations will yield poorly defined shape graphs. In the following sections, we will present several possibilities of parameters one could choose for each Mapper step. The parameter choices and their impact on the final shape graph will be presented as empirical results.

#### 2.1.1 Distance metric

The first step of the Mapper algorithm is defining a distance metric for the dataset, designating the relationship between points in the original high-dimensional space. The distance metric picked is the main parameter defining this step. Here we analyzed three broad measures of distance: angle-based measures (Cosine and Correlation), magnitude measures (Euclidean, Cityblock, and Chebychev) (Bobadilla-Suarez et al. 2020), and the geodesic (or shortest path) metric. We use the following metric definitions for the two vectors *x* and *y*, representing the distance between those two points:

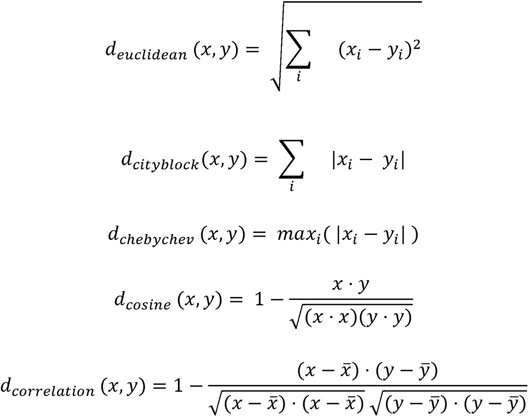

Where

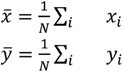

And the operation *a* · *b* is the dot product between vectors *a* and *b*.

For the geodesic metric, we first constructed a penalized reciprocal k-nearest neighbors graph (PRKNNG), then considered the distance between points as hops on this constructed graph. The reciprocal variant of the k-NN algorithm limits to only connections between points that the k-NN bidirectionally links, thereby reducing the effect of outliers (Qin et al. 2011). Additionally, to fully connect the resulting penalized k-NN graph, we added back connections between clusters with exponentially penalized weights (Bayá and Granitto 2011). We showed in previous work (Geniesse, Chowdhury, and Saggar 2022) that the reciprocal and penalized variant of the k-NN algorithm works synergistically with Mapper. It’s important to note that this algorithm requires a distance metric (e.g., Euclidean, Cosine) to calculate the k-NN graph:

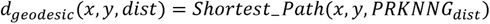

Where

*PRKNNG_dist_* is the Penalized, Reciprocal k-Nearest Neighbor graph with weighted edges constructed using the distance metric *dist*

Defining the metric space is essential for the partial clustering step where we evaluate distances between the data points in the original high-dimensional space (Singh et al. 2007). Moreover, some filter functions use distances between all pairs of data points (pairwise distances). Due to the high computational cost of computing certain pairwise distances, we did not use distributional magnitude measurements like Mahalanobis and Bhattacharyya distances.

#### 2.1.2 Filtering

During the second step of the Mapper algorithm, the data points are projected, using a filter function, to a reduced space (**Fig. 1c**). The filter function can be applied to either the pairwise distances or the original data space. Possible filter functions (aka lenses) include dimensionality reduction techniques (**Fig. 1c top**) that have been previously explored and analyzed within the neuroscience field (Cunningham and Yu 2014). The data is usually reduced to a few (2 or 3) dimensions for practical and visualization purposes as the binning step scales exponentially with the number of dimensions. Any dimensionality reduction method can be used as a filter, but some have desirable properties and can better preserve the topological features of the dataset. In this work, we compared multiple types of dimensionality reduction algorithms (**Table 1**). Because some selected algorithms (UMAP, Isomap, LLE, HessianLLE, Laplacian, and LTSA) construct k-NN maps and pairwise distances as a step, we applied them directly to the original dataset.

**Table 1:** Dimensionality reduction algorithms. The ‘Code’ column represents the short-hand form used for the rest of the paper. The ‘Applied on’ column represents the usage of each algorithm as they were applied on either the pairwise distances or the original dataset space.

| Name | Code | Applied on (Input) | Reference |
| --- | --- | --- | --- |
| Classical Multidimensional Scaling | CMDS | Pairwise distances | (Seber 2009) |
| Principal Component Analysis | PCA | Pairwise distances | (Pearson 1901) |
| Linear Discriminant Analysis | LDA | Pairwise distances | (Fisher 1936) |
| Factor Analysis | FactorAnalysis | Pairwise | (Spearman 1904) |
|  |  | distances |  |
| Diffusion Maps | DiffusionMaps | Pairwise distances | (Lafon and Lee 2006) |
| Sammon mapping | Sammon | Pairwise distances | (Sammon 1969) |
| Uniform Manifold Approximation and Projection | UMAP | Original data | (McInnes, Healy, and Melville 2018) |
| Isomap | Isomap | Original data | (Tenenbaum, de Silva, and Langford 2000) |
| Locally Linear Embedding | LLE | Original data | (Roweis and Saul 2000) |
| Hessian Locally Linear Embedding | HessianLLE | Original data | (Donoho and Grimes 2003) |
| Laplacian Eigenmaps | Laplacian | Original data | (Belkin and Niyogi 2001) |
| Local Tangent Space Alignment | LTSA | Original data | (Z. Zhang and Zha 2004) |
| t-distributed Stochastic Neighborhood Estimation | t-SNE | Pairwise distances | (Hinton and Roweis 2002) |

While there are even more options for dimensionality reduction, we focused on the main algorithms used with Mapper in the literature in this paper. We point the reader to a review for a comprehensive analysis and a systematic comparison of dimensionality reduction techniques (Van Der Maaten et al. 2009).

To avoid dimensionality reduction and the related information loss altogether, our group recently developed a filter function that operates directly on the penalized k-NN graph constructed in the original high-dimensional space (**Fig. 1c bottom**) (Geniesse, Chowdhury, and Saggar 2022). For this technique, the lens is simply the geodesic distances constructed in the previous step, maintaining the local structure for each data point. As the locality is preserved, we define a Mapper that uses this technique as an **Intrinsic Mapper**. An **Extrinsic Mapper**, on the other hand, uses a dimensionality reduction technique as a filter function. As the two types of lenses, i.e., intrinsic vs. extrinsic, operate in different spaces, each requires its own binning algorithm. Even though the two mapper types diverge during the filtering and binning steps (**Fig. 1c-d arrows**), they can use the same partial clustering and graph construction method.

#### 2.1.3 Binning

The third step of Mapper consists of segmenting the data into overlapping bins. For the Extrinsic Mapper, the data contains points in a low-dimensional space, and the binning consists of segregating the points into overlapping bins (**Fig. 1d top**). The bins could be any shape that collectively covers the whole embedding space. For 2-dimensional embeddings, the bins are polygons, with rectangles being the most commonly used. Each bin size and placement are determined by the scale parameters of this step: the resolution and gain. The resolution is the number of bins per dimension, and the gain is the amount of overlap between the bins, measured in percentages.

For the Intrinsic Mapper, binning requires segmenting the constructed k-NN graph into overlapping bins (**Fig. 1d bottom**). In this case, the bins represent N-dimensional spheres centered around specific landmarks picked from data points using the Farthest Point Sampling (FPS) algorithm (Geniesse, Chowdhury, and Saggar 2022). Each bin size and location are determined by the scale parameters: the resolution and gain. The resolution represents the number of landmarks chosen, while the gain defines the distance around each landmark to include within a bin (**Fig. 1d bottom**). Specifically, a datapoint *x_i_* is considered within a bin minimum distance between any two landmark *x*, if 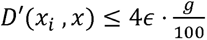, where D’ is the distance metric used, 2*ɛ* is the minimum distance between any two landmarks, and *g* is the gain parameter. The gain parameter approximates the overlap between the spherical bins, and the values are picked from 0 to 100, representing a percentage. Each generated bin will contain a set of data points that will be further clustered in the following steps of the Mapper algorithm.

While both binning steps (extrinsic and intrinsic) are parameterized by resolution and gain, the parameters’ values have different connotations and might result in qualitatively different shape graphs. For this reason, a direct comparison between the shape graphs resulting from the two methods is rendered incompatible. Instead, in our analysis, we contrast the data point connectivity matrices that result from the Mappers of those binning strategies (Extrinsic Mapper vs. Intrinsic Mapper). An in-depth mathematical justification for the Intrinsic Mapper as a valid topological tool was performed in our previous paper (Geniesse, Chowdhury, and Saggar 2022). In this work, we provide further evidence of the equivalence of the two mappers based on their generated connectivity matrices over large spaces of parameter configurations.

#### 2.1.4 Partial Clustering

Once the data points are assigned to bins, the fourth step of the Mapper algorithm involves clustering those data points (**Fig. 1e**). Importantly, the clustering algorithm is run for each bin independently, and clustering is done on the original feature space (Singh et al. 2007). The generated clusters constitute the nodes of the shape graph.

The Mapper algorithm uses any hierarchical or nonhierarchical clustering technique (James et al. 2013), such as single linkage (Landau et al. 2011), average linkage, or density-based spatial clustering of applications with noise (DBSCAN) (Ester et al. 1996).

#### 2.1.5 Graph Creation

As a final step, the Mapper algorithm links (with an edge) the created nodes that share at least one data point. This step is made possible because the binning step provides a degree of overlap (gain), representing data points in multiple bins. The nodes and edges comprise the Mapper “shape graph,” representing the topology of the input dataset. The resulting shape graph is an undirected graph as the edges are bidirectional.

The graph creation step does not require any parameters. Although, one alternative to the graph construction step is to limit the edges to bins that are adjacent to each other (van Veen et al. 2019). For example, when using a 2-dimensional embedding with rectangle bins, a node will be limited to the eight directly adjacent bins, even though there might be more overlapping bins (when *gain* > 50%). This variation can only be performed in Extrinsic Mapper settings, and the shape graph produces a grid-like pattern.

Another alternative is constructing a directed shape graph based on the temporal progression of the data points (M. Zhang, Chowdhury, and Saggar 2022). This “Temporal Mapper” requires using a different filter function and a modified binning step, but this work did not analyze its application.

### 2.2 Datasets

#### 2.2.1 Dataset 1: Simulated temporal structure using a biophysical network model

To generate gold standard ground truth transitions of brain activity, we used a dynamical mean-field model of a human brain to generate controlled BOLD signals. Complete details of the model and how these synthetic data were generated are presented elsewhere (M. Zhang, Sun, and Saggar 2022). Briefly described here, we used a biophysical network model that adapted the reduced Wong-Wang (Deco et al. 2013, 2014; Wong and Wang 2006) model in the form of Wilson-Cowan model (Wilson and Cowan 1972, 1973) to improve multistability (M. Zhang, Sun, and Saggar 2022). The model constructs a large-scale network (global model, **Fig. 2b**) using nodes corresponding to anatomical regions in the human brain based on 66-region parcels (Deco et al. 2013). The network’s edge weights between the associated brain regions are estimated using structural connectivity data from the Human Connectome Project (Van Essen et al. 2013). Each node is modeled as a pair of excitatory (E) and inhibitory (I) populations with four connections describing their influence: *W_EE_* modulating *E* population exciting itself *W_EI_* modulating *E* population exciting the *I* population; *W_II_* modulating *I* population inhibiting itself; *W_IE_* modulating *I* population inhibiting the *E* population (local model) (**Fig. 2a**). The state variables *S_E_* and *S_I_* describe the activity of the two populations within each node, and physically, they represent the fraction of open synaptic channels in each population. The long-range connections of the global model are between the excitatory populations of each node and are modeled by the static variable *C_ij_*. Furthermore, the overall strength of those long-range connections is scaled by a global coupling parameter *G*.

**Figure 2:**
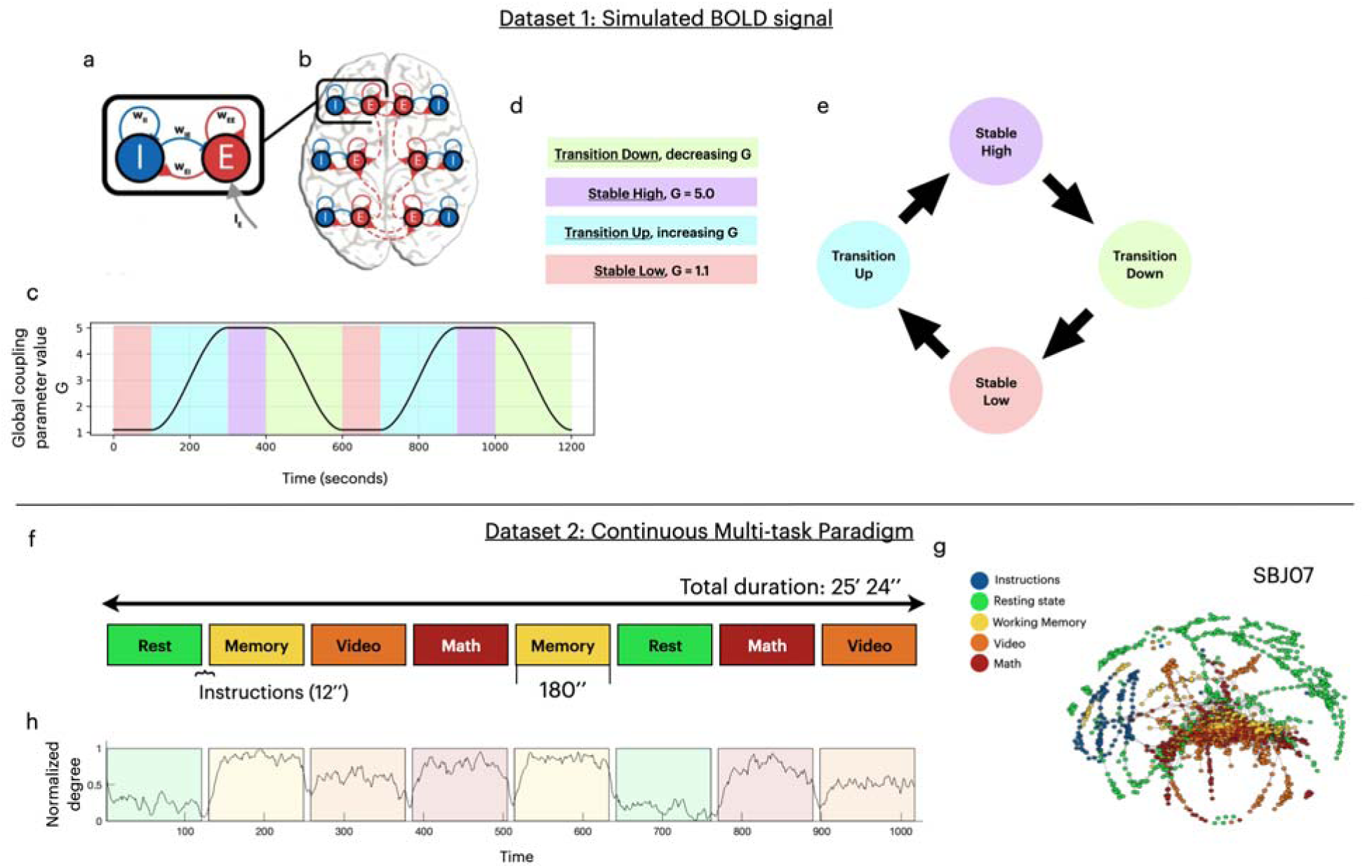
Datasets description. (a-e) Dataset 1: Simulated BOLD. **(a)** The local model connectivity between the pair of populations, excitatory (E) and inhibitory (I), is defined by four connections:, and. **(b)** The global model defines the connectivity profile between the nodes, connecting the excitatory populations. **(c)** The timeline of values of the global coupling parameter value, throughout the course of the simulation. **(d)** The selected four states of the simulation time course. **(e)** A transition graph representation between the four states. **(f-h) Dataset 2: Continuous Multi-Task Paradigm (CMP), the “real” dataset. (f)** Representation of the timeline of four tasks in the CMP dataset: Resting state, Working Memory, Video, and Math. Each task was performed for 180 minutes with 12 seconds of instruction in between. The total duration of the scan was 25 minutes and 24 seconds. **(g)** Example of a constructed Mapper shape graph on subject SBJ07 using an Extrinsic Mapper (geodesic Euclidean metric, k=12, CMDS embedding, resolution=30, gain=60%, Linkage clustering). **(h)** The normalized degree, averaged over subjects, was extracted from the Mapper shape graphs.

The global coupling parameter was modulated between two values to create “true” temporal variation in the generated neural activity (**Fig. 2c**). We can separate the time course of the simulation into four types of states (**Fig. 2d**) based on the value of the parameter. The Stable Low and Stable High states are the time points where is stable at either a low or high value, respectively. In contrast, the transition between the two stable states is considered the Transition Up or Transition Down states (**Fig. 2d**). The transitions between the four states and the expected dynamical topology *represent a circle with a preferred direction* (**Fig. 2e**). To generate the BOLD signal, the neural activity of each modeled brain region, represented by the local excitatory population activity, was fed into the traditional Balloon-Windkessel model (Buxton, Wong, and Frank 1998). Using a Repetition Time of 0.72 seconds, we generated 20 minutes (1200 seconds) of data, totaling 1667 data points.

We extended this synthetic dataset by adding noise and downsampling to test the Mapper algorithm under higher noise regimes and with lower temporal resolution acquisitions. For adding noise to our signal, we first generated white noise using the ‘brainiak’ package (Ellis et al. 2020) based on the extracted properties of the input dataset: drift noise, auto-regressive/moving-average noise, and system noise. Then, based on the desired Signal-To-Noise ratio (SNR), we scaled and added the generated noise to each voxel activation of the dataset. Similarly, we extended the original synthetic (without the added noise) by downsampling (same as increasing repetition time) to test the effect of lowering the temporal resolution of the signal. First, we temporally smooth the data to allow the information to be preserved over the remaining data points. Then, we selected every other (TR doubles) or every third (TR triples) data point, thus creating the new extended datasets.

#### 2.2.2 Dataset 2: Real temporal structure using CMP Dataset

As our second dataset, we used a previously collected dataset by (Gonzalez-Castillo et al. 2019). The dataset uses a Continuous Multitask Paradigm (CMP), scanning participants while performing an array of tasks (**Fig. 2f**). We transferred the dataset from the XNAT Central public repository (https://central.xnat.org; Project ID: FCStateClassif). All participants provided informed consent, and the local Institutional Review Board of the National Institute of Mental Health in Bethesda, MD, reviewed and approved the data collection.

The CMP dataset contains de-identified fMRI scans with their associated behavioral measurement from 18 participants. The complete details of the paradigm are presented in (Gonzalez-Castillo et al. 2019). As described here briefly, the participants performed four different tasks, each repeated once, while being scanned continuously inside an MRI machine. The four types of tasks were classified as Rest, Working Memory, Math/Arithmetic, and Video; each being carried out for 180 seconds, with an extra 12-second instruction period. As each task was repeated, the final eight task blocks appeared in a predetermined random order, similar for all participants. During the Rest task, each participant was instructed to fixate on a crosshair at the center of the screen and let their mind wander. For the Working Memory task, the participants were presented with geometric shapes every 3 seconds and were instructed to signal (by pressing a button) if the current shape appeared two shapes prior (2-back design). For the Math/Arithmetic task, the participants were sequentially presented with 36 total arithmetic operations, while each one involved applying two operators (addition and subtraction) on three numbers between 1 and 10. During the Video task, the participants watched a video of a fish tank from a single stationary point of view with different types of fish swimming into and out of the frame; the participants were instructed to press a button when a red crosshair appeared on a clown fish and another when it appeared on any other type of fish.

The fMRI dataset was acquired on a Siemens 7 Tesla MRI scanner equipped with a 32-channel receiver coil (Nova Medical) using a whole-brain echo planar imaging (EPI) sequence (repetition time [TR] = 1.5 s, echo time [TE] = 25 ms, and voxel size = isotropic 2 mm). A total of 1017 time frames were acquired for each participant.

Functional and anatomical MR images were preprocessed using the Configurable Pipeline for the Analysis of Connectomes (C-PAC version 0.3.4; https://fcp-indi.github.io/docs/user/index.html). Complete details about the processing are provided by (Saggar et al. 2018). Briefly, both anatomical and functional scans were registered into the MNI152 space (using ANTS) after registering each participant’s functional scan to match its corresponding anatomical scan. Further, the fMRI data preprocessing steps included slice timing correction, motion correction (using the FSL MCFLIRT tool), skull stripping (using the FSL BET tool), grand mean scaling, spatial smoothing (FWHM of 4mm), and temporal band-pass filtering (between 0.009 Hz and 0.08 Hz). Additionally, nuisance signal correction was done on the data by regressing out (1) linear and quadratic trends; (2) physiological noise (mean time-series of white matter and cerebrospinal fluid); (3) derived motion noise from 24 motion parameters (the six motion parameters, their derivatives, plus each of these values squared); and (4) signals extracted using the CompCor algorithm (five selected components). Finally, the resulting voxels were averaged to 3 mm MNI space and further fit within the 375 regions of interest (ROIs) with 333 cortical parcels (Gordon et al. 2016) and 42 sub-cortical parcels from the Harvard-Oxford atlas (Shine et al. 2016).

### 2.3 Mapper testing

To examine Mapper parameters and the quality of the final Mapper graph, we devised general criteria for shape graphs validation and goodness-of-fit measures (GOF) for each dataset. These GOF measures take into account the expected properties of each dataset.

#### 2.3.1. Validating shape graphs

Drawing on prior knowledge and validated expectations of valid shape graphs (Geniesse, Chowdhury, and Saggar 2022), we developed three metrics that validates the coverage, autocorrelation, and complexity captured. We test if the Mapper shape graph: (i) captures the shape of most of the input dataset (coverage β > 70%); (ii) captures more than the autocorrelation dynamics (non-autocorrelated nodes *α* ≥ 15%); (iii) has a non-trivial structure (pairwise distances entropy *S* ≥ 2). We define the Mapper shape graph coverage (*β*) as the percentage of data points in the largest connected component of the shape graph. To measure the influence of autocorrelation dynamics, we count the percentage of nodes (*α*) that describe data points over the autocorrelation time threshold, *τ*. We chose *τ = 11S*, as its generally expected to be the hemodynamic response function peak for the BOLD neural response (Lindquist et al. 2009). In addition to the properties described in the previous study by Geniesse et al. (2022), we introduce a novel metric to remove degenerate shape graphs. We observed that for some Mapper configurations, the shape graph nodes connect into large cliques, destroying all topological properties of the input dataset. Hence, we quantify and invalidate the shape graphs that have a low entropy (*S*) (Shannon 2001) of pair-wise distances between all nodes of the graph.

#### 2.3.2 Goodness of fit measures for the Simulated Dataset

A biophysical network model was used to generate simulated transitions in whole-brain activity. These transitions were generated with a true shape of topology that *represented a circle with a preferred direction.* Thus, to quantify if the resulting shape graph correctly represents the expected transition graph for the simulated data, we defined a “circle-ness” criterion as a GOF measure (**Fig. 2e**). A good quality Mapper shape graph should contain nodes that connect the low and high states bidirectionally by edge paths through the two transition states, respectively. The algorithm to test if a Mapper shape graph satisfies the circleness criterion is explained as follows. First, we mark each node as one of four states: stable-low, transition-up, stable-high, and transition-down, based on the states of the data points it represents. Then we create the graph *G*_↑_ as a subset of the shape graph, excluding the nodes marked as state transition-down. We test if the graph *G*_↑_ contains a path between nodes describing stable-low and stable-high states, using only nodes that are transition-up. Similarly, we test the sub graph *G*_↓_ (the shape graph without the transition-up nodes) if it contains the same path but now using only nodes that are transition-down. If the shape graph passes both test, then it is passes the circleness criterion.

#### 2.3.3 Goodness of fit measures for the Real Dataset

To quantify the quality of the Mapper graphs generated from the real neuroimaging dataset, we constructed a GOF metric that examined the transition times between four cognitive tasks. This metric is the average delay time, which measures the time difference between predicted and expected state changes, delimiting the four states (Rest, Memory, Video, and Math). We have previously shown that by extracting the similarity between individual time frames from the Mapper graph, we can track transitions in brain states at the individual time frame level (Saggar et al. 2018). Thus, time frames that belong to the same node or are connected by an edge in the Mapper graph are considered highly similar. Estimating similarity between time frames can lead to a temporal connectivity matrix (TCM), which shows similarity between each time frame to every other time frame in the session. Using the average temporal similarity of time frames, referred to as normalized degree of the TCM, we extracted the predicted state changes for all participants, where abrupt changes in the similarity of time frames indicated a state change (**Fig. 2h**). To find the abrupt changes in the normalized degree timeline, we employ a Changepoint Detection algorithm, implemented in MATLAB by the function *findchangepts* (Killick, Fearnhead, and Eckley 2012). Having the average delay time of a Mapper result allowed us to compare different parameter configurations to find the most suitable parameters.

### 2.4 Data and code availability

The synthetic datasets used in this work and all the associated code will be available upon publication at this address: https://github.com/braindynamicslab/demapper. The fMRI data used in this study is available for download at the XNAT Central public repository (https://central.xnat.org; Project ID: FCStateClassif).

## 3. Results

### 3.1 Similarity between individual time frames

The first step of the Mapper algorithm is computing pairwise distances between the input data points. While this is a straightforward computational task, choosing a distance metric has wide implications because it defines the relationship between any two points for the rest of the algorithm. Finding a correct similarity metric (i.e., *similarity* = 1 − *distance*) samples of neural activity is a long-studied topic in neuroscience (Bobadilla-Suarez et al. 2020). The goal of this paper is not to solve the issue but rather to reveal the effects of choosing different distance metrics for the Mapper algorithm. Here we analyzed three broad measures of distances: angle-based measures (Cosine and Correlation), magnitude measures (Euclidean, Cityblock, and Chebychev) (Bobadilla-Suarez et al. 2020), and geodesic metrics.

We used the “circleness criterion” as a goodness of fit (GOF) metric for the simulated dataset. While varying the distance metrics, we examined several combinations of other Mapper parameters (e.g., resolution and gain). **Fig. 3** shows a pattern of Mapper results that satisfy the criteria. As an example, with a correlation distance metric, the resulting shape graph created high similarity between transition-up and transition-down states (**Fig. 3a**). This degeneration of the expected connectivity between states leads to the rejection of this shape graph as a correct topological representation of the input (circleness, see methods). For a selection of different resolution and gain parameters, the geodesic Euclidean distance metric (with k=12) yields 19 out of 25 graphs that preserve the expected features (**Fig. 3b center**). We assessed the shape graphs of using other distance metrics on the same grid of resolution and gain parameters (**Fig. 3b**). Alternating the k-parameters for the geodesic distances, the different configurations of the distance metrics yield a distribution of Mapper shape graphs that pass the criterion (**Fig. 3c**). Choosing a distance metric has a significant impact on the performance of the topological extraction (ANOVA one-way, p < 10^-5^). Specifically, the Euclidean and Cosine geodesic distance metrics generally perform better (**Fig. 3c**). Furthermore, we observe no significant difference between the performance of magnitude and angle metrics (t-test, p = 0.9).

**Figure 3:**
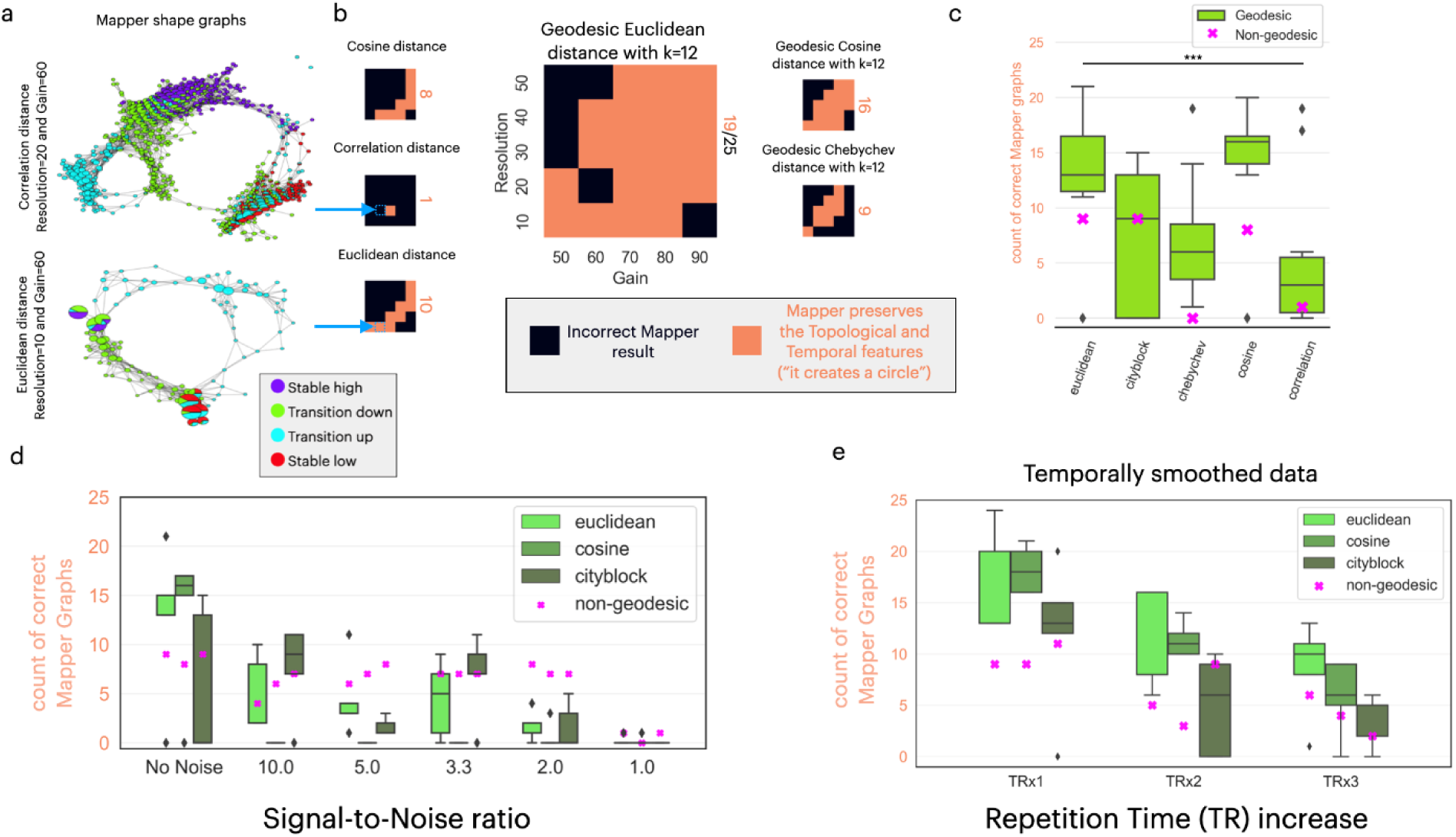
Quantifying similarity metrics performance on the simulated dataset. **(a)** Example of two Mapper shape graph results. **Top**: using the correlation distance (resolution=20, gain=60%). This shape graph is classified as an “incorrect Mapper result” because the “transition up” nodes do not define a path between nodes of stable up and down states. **Bottom**: using the Euclidean distance (resolution=10, gain=60%). This shape graph is valid as it passes the Mapper shape graph validity metrics and GOF metrics for the simulated dataset. **(b)** The performances of different distance metrics are shown as a heatmap on a resolution-by-gain 5×5 matrix. Each resolution-gain parameter choice within the heatmap represents the Mapper algorithm’s success in preserving the circular state trajectory’s topological and temporal features. The total count of such correct Mapper results is presented as an orange letter on the right of each heatmap. The two examples in **(a)** have two squares in their respective heatmaps, defining their validity. **(c)** The five distance metrics show different performances in capturing the expected circle trajectory, with the Euclidean and Cosine geodesic distances outperforming the rest. The non-geodesic distances are represented as a purple “X” marker for each distance metric. The line with *** denotes a one-way ANOVA with p < 10^-5^. **(d)** As we decrease the signal-to-noise ratio, the performance decreases for all distance metrics, with the Cosine distance decreasing the most, revealing a sensitivity to noise. **(e)** With a decrease in the sampling rate of the simulated dataset, we see a decrease in the performance for all distance metrics. The distributions in subplots (c), (d), and (e) are shown as box plots.

We observe that the average performance of using geodesic distances (averaged over k-values) is higher than its relative non-geodesic distance performance, but it fails the significance test due to a low sampling rate (t-test relative, p = 0.08). On this dataset, we observe that the geodesic distance metrics require a minimal k-value: Euclidean and Cosine distance metrics require *k* ≥ 6; while the Cityblock distance metric requires *k* ≥ 32 (Supplementary **Fig. S1**). Similar results were observed for the intrinsic mapper when using a k-NN lens instead of reducing the embedding space using a dimensionality reduction technique (Supplementary **Fig. S1**).

We also evaluated the effect of increasing noise in the data by analyzing the top distance metrics (Euclidean, Cosine, and City-block) on the simulated dataset with decreasing signal-to-noise ratio (SNR). We constructed this noisy dataset by progressively adding white noise to all regions of the simulated dataset. As expected, we observe a general decrease in performance as we decrease the SNR. The rate of decrease is more pronounced in geodesic distance metrics compared to non-geodesic metrics (t-test relative for selected distances, p < 0.005) (**Fig. 3d**, Supplementary **Fig. S2a**). This suggests that non-geodesic distances are more robust to white noise in this simulated dataset. Moreover, the geodesic angle metrics (cosine and correlation) fail to construct valid mapper graphs once we introduce noise (Supplementary **Fig. S2a**).

Further, we evaluated the performance of the Mapper shape graph on the simulated dataset with a decreased sampling rate (or an increased repetition time). We constructed this dataset by temporally smoothing the data and sampling every second (TRx2) or every third (TRx3) data point. We observe that the performance decreases as we decrease the sampling rate across all distance metrics, showing no difference between them (**Fig. 3e**). As seen in the general case, the average performance of using geodesic distances (averaged over k-values) is higher than its relative non-geodesic distance performance (t-test relative, p < 0.001). This relative performance improvement when using geodesic distances is observed in all metric spaces (correlation and chebychev metrics are shown in Supplementary **Fig. S2c**).

We analyzed the distance metrics for the real dataset for each shape graph based on the average delay between the expected and predicted transitions (see Methods). For example, using the geodesic Euclidean distance (with k=12), Mapper predicted the transitions between the eight states with an average delay of 5.7 seconds (**Fig. 4a top row**). In contrast, using the Euclidean distance (non-geodesic), Mapper failed to predict the 7^th^ transition between the Math and Video states (**Fig. 4a bottom row**). Aggregating over multiple shape graphs, the choice of distance metrics has a significant impact on the performance of the topological extraction (ANOVA one-way, p < 10^-10^) (**Fig. 4b**). Moreover, the magnitude metrics outperform the angle metrics (t-test, p < 10^-10^), without a clear magnitude metric performing best (ANOVA one-way, p > 0.05) (**Fig. 4b**). Furthermore, the average performance of using geodesic distances (averaged over k-values) is higher than its relative non-geodesic distance performances (t-test relative, p < 0.05). Those three findings and tests are replicated when we use a higher delay threshold of 20 seconds (Supplementary **Fig. S2d**).

**Figure 4:**
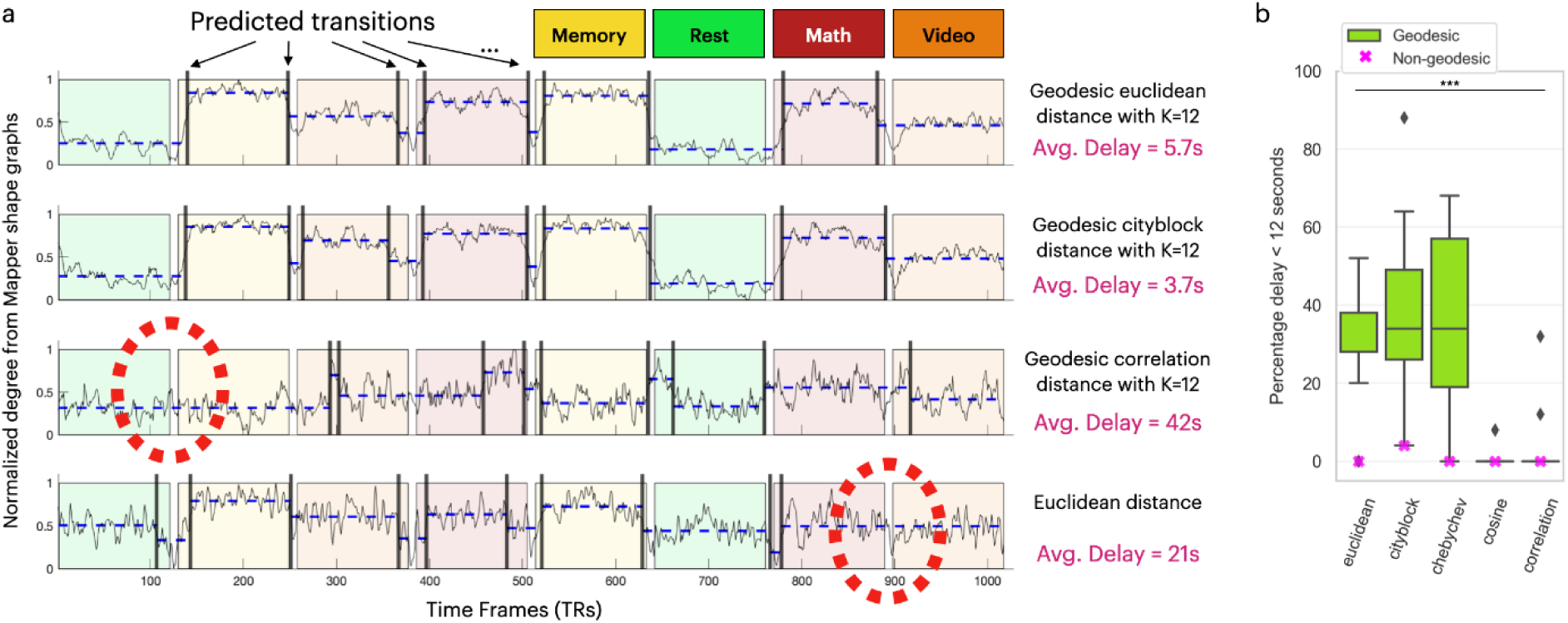
Quantifying similarity metrics performance on the real dataset. **(a)** Four examples are presented based on different distance metrics for generating the Mapper graph. The examples are for Mapper Graphs generated with Resolution=20 and Gain=50. The x-axis timeline is divided into eight regions, colored based on the task performed during that time. The timeseries shown as a light black line is the normalized degree of the Mapper shape graph. The timeseries was used to extract predicted transitions, marked as vertical black lines. The dashed blue lines between the predicted transitions represent the level of average normalized degree. The large red circles with dashed lines highlight regions that failed to be predicted as a transition. The match between expected and predicted is quantified as an average delay. The first two examples have a short delay (geodesic Euclidean distance with K=12: 5.7 seconds; geodesic Cityblock distance with K=12: 3.7 seconds), while the last two examples have a large delay due to the missed predictions (geodesic Correlation distance with K=12: 42 seconds; non-geodesic Euclidean distance: 21 seconds). **(b)** For multiple values of resolution, gain, and K-values, the performance of different distance metrics is shown as a percentage of average delays smaller than 12 seconds. The geodesic distributions are shown as boxplots. The non-geodesic distances are represented as a purple “X” marker for each distance metric. The line with *** denotes a one-way ANOVA with p < 10^-10^.

### 3.2 Effect of the embedding algorithm on the Mapper results

The traditional Mapper algorithm (Extrinsic Mapper) represents data points in a lower dimensional space. This embedding is often created using dimensionality reduction techniques with different assumptions about the represented topology and the relevant features. We measured the performance of several embedding algorithms on the simulated and real neuroimaging datasets. For each embedding algorithm, we count the mappers that fulfill the GOF criteria with different k-values for either the geodesic distance metric or the embedding algorithm (**Fig. 5**). For the simulated dataset, we observe that multiple algorithms (CMDS, PCA, LDA, FactorAnalysis, Sammon, Isomap) perform almost identically (**Fig. 5a**). While UMAP has low performance, we see that it requires low values of the k-parameter (i.e.,), after which the performance drops to zero (**Fig. 5b**). Although, this inconsistency of the k-parameter is due to the UMAP algorithm performing its topological deconstruction. Moreover, the t-SNE embeddings fail to extract the topological features for the simulated dataset (**Fig. 5a**). Examples of the created shape graphs using t-SNE demonstrate that while the local structure is preserved, it fails to construct the whole circle (Supplementary **Fig. S4**).

**Figure 5:**
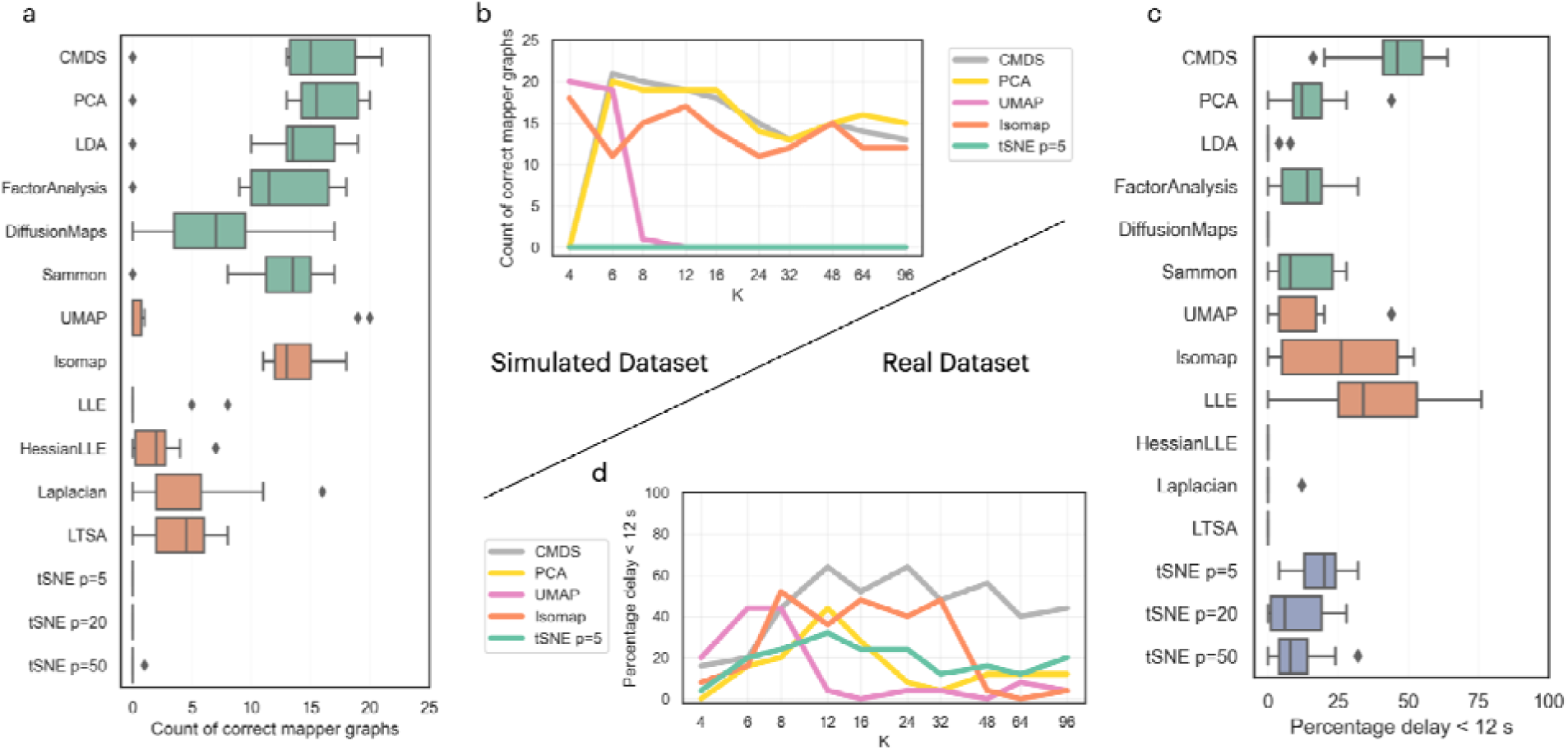
The effect of the embedding algorithm choices for constructing the Mapper Shape graphs. **(a)** Performance of Mapper on the simulated dataset using different embedding algorithms, where each box plot corresponds to the distribution of shape graphs that pass the GOF criterion (based on different k-values). The top six box plots (denoted in green) represent dimensionality reduction techniques applied on pairwise distances (CMDS, PCA, LDA, FactorAnalysis, DiffusionMaps, Sammon), with the distribution based on the geodesic k-value used for the distances. The following six box plots (denoted in orange) represent dimensionality reduction techniques applied on the original space (UMAP, Isomap, LLE, HessianLLE, Laplacian, LTSA), with the distribution based on the k-value used for applying the embedding algorithm on the input dataset (skipping the pairwise distances). The bottom three box plots (denoted in purple) represent the distribution of performance of the stochastic algorithm (t-SNE) using different perplexity values (5, 20, 50), with the distribution of a geodesic k-value. **(b)** A few selected algorithms’ performance was broken down based on different k-values on the simulated dataset. **(c)** The performance of Mapper on the real dataset using the same embedding algorithms as subplot (a). **(d)** The same selected algorithms’ performance is broken down on a set of k-values on the real dataset. The performance distributions in (a) and (c) are shown as box plots.

On the real dataset, the CMDS embedding algorithm constructs better representations than the other embedding algorithms applied on pairwise inputs (**Fig. 5c**). Comparing the non-pairwise algorithms, we see Locally Linear Embeddings (LLE) and Isomap having better representations. As seen in the simulated dataset, the UMAP algorithm requires lower values of the k-parameter to construct good representations (**Fig. 5d**). In this case, the t-SNE algorithm has more success in creating shape graphs that pass the validation criterion.

An alternative to embedding algorithms is the intrinsic mapper algorithm, which performs the topological analysis in the original space (Geniesse, Chowdhury, and Saggar 2022). While the Intrinsic Mapper algorithm has different parameters (resolution represents the number of landmarks instead of the number of bins), it generates remarkably similar Mapper shape graphs (Supplementary **Fig. S5**). The extrinsic and intrinsic mappers produce similar distances between time frames, as measured by their corresponding temporal connectivity matrices (TCMs) (Supplementary **Fig. S6**). Moreover, the intrinsic mapper projects the data to a space that resembles a high dimensionality embedding (Geniesse, Chowdhury, and Saggar 2022), which would not be achievable with extrinsic mapper because of the exponential explosion of bins (i.e., for resolution R and dimensions d, we have R^d^ bins). Hence, the intrinsic mapper allows for faster processing of bins/landmarks for clustering and creating the shape graph nodes as we have increasingly more nodes (Supplementary **Fig. S6**).

### 3.3 The appropriate scale of reduction for neuroimaging data

The Mapper graph attempts to reveal the shape of the high-dimensional input data in a low-dimensional space. As for any algorithm that compresses information, the representation can be underfitting or overfitting the data. In the context of a topological analysis, we expect the representation to preserve the topological features with the right amount of detail. The resolution and gain parameters during the binning step of the Mapper algorithm determine the size of the shape graph (**Fig. 1d**). Selecting the appropriate scale of reduction (i.e., resolution and gain) is a necessary step for configuring Mapper to extract the topological and temporal features of any time-series dataset. Different scale parameters (i.e., resolution and gain) can result in qualitatively different shape graphs (**Fig. 6a-c**).

**Figure 6:**
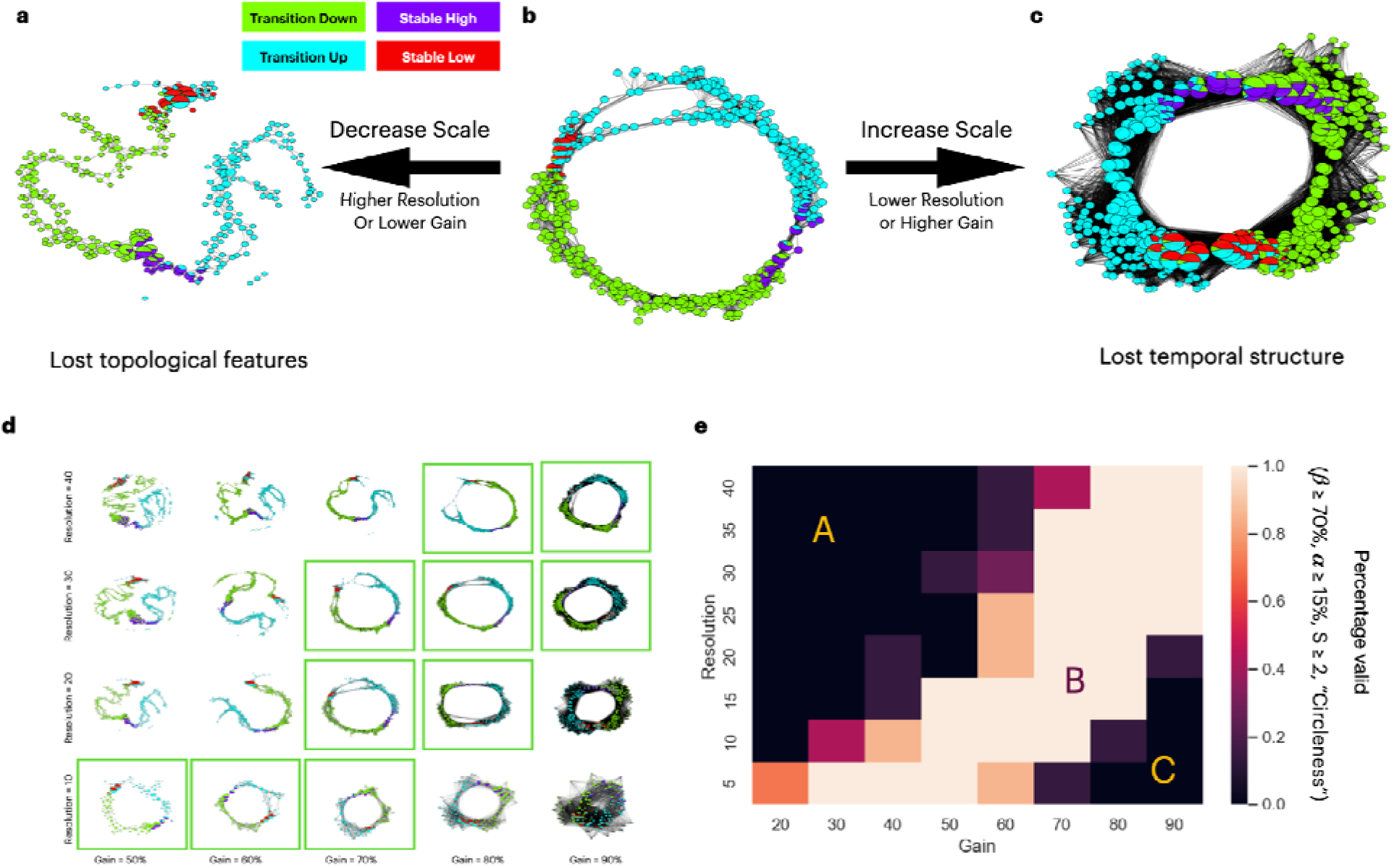
Choosing the appropriate scale when running Mapper. **(a)** Shape graph results from the simulated dataset produced by an Extrinsic Mapper with resolution=20, gain=50%, k=20. Each node is represented as a pie chart of the composition of time points within that node. **(b)** Shape graph result using Extrinsic Mapper with resolution=20, gain=70%, k=20. **(c)** Shape graph result using Extrinsic Mapper with resolution=20, gain=90%, k=20. **(d)** Grid of shape graph produced by Extrinsic Mappers with k=20 for resolution parameters: 10, 20, 30, and 40; and gain parameters: 50%, 60%, 70%, 80%, and 90%. The valid Mapper results are highlighted within a green box. **(e)** A larger grid of valid Mapper results is now aggregated over different k values: 10, 20, 30, 40, 50, 60, and 70. The plot shows three main regions. Region A is associated with high resolution and/or low gain, similar to subplot (a). Region C is associated with low resolution and/or high gain, similar to subplot (c). The middle region B shows a band of valid Mapper graphs independent of any k value with appropriate resolution and gain parameters. All Mappers in this plot are generated with the geodesic Euclidean distance metric (needs the k parameter), CMDS embedding, extrinsic binning, and single linkage clustering.

Starting with the simulated data, Mapper graphs with low gain (Extrinsic Mapper with resolution 20, gain 50%) do not capture the circular pattern of the neural input data (**Fig. 6a**). This failure is due to the discontinuity within the *transition-up* timeframes. Because of the missing edges in the result, the topological feature of the input dataset was not preserved, and we mark this result as a failure of the parameter choices. In contrast, the circular topological feature is correctly revealed (**Fig. 6b**) by the Mapper configuration with an increased gain value (Extrinsic Mapper with resolution 20, gain 70%). This combination of resolution and gain creates a shape graph that represents the correct transition between the time points as it was originally generated. Moreover, a Mapper with an even higher gain (Extrinsic Mapper with resolution 20, gain 90%) creates a highly connected graph that directly links the stable-low and stable-high states (**Fig. 6c**). This high connectivity loses the specificity of the topological structure by bypassing the temporal profile of individual timeframes. As we mark this result as a failure, we can now intuitively appreciate the boundary of parameter combinations. We reveal a distribution of valid shape graphs where the resolution and gain parameters are highly correlated (**Fig. 6d**). Aggregating on multiple k-values (for the geodesic distance) results in a similar correlation between resolution and gain (**Fig. 6e**). For high resolution or low gain, the shape graphs lose the topological features, showing a discontinuity between stable-low and transition-up states (**Fig. 6e section A, Fig. 6a**). On the other side, for high gain or low resolution, the result loses the temporal structure and fully connects the shape graph (**Fig. 6e section C, Fig. 6c**). In the middle, for adequate combinations of resolution and gain, the topology of the input dataset is preserved and correctly represented by the shape graph (**Fig. 6e section B, Fig. 6b**). This distribution of valid results based on scale parameters provides guidance on choosing an appropriate combination. Proportional changes in gain and resolution parameters will yield the same topological features. Moreover, increasing the scale parameters will increase the total number of nodes, with the resolution parameter having a greater effect (Pearson r=0.8687, p<10^-10^) than the gain parameter (Pearson r=0.3617, p<10^-10^).

Using a different binning strategy, we observe the same parameter dependence (Intrinsic Binning, see Methods), where the resolution parameter controls the number of landmarks chosen on the k-NN graph, and the gain controls the distances and overlap between landmarks (Supplementary **Fig. S5**). As the filtering function might influence the types of topological features extracted, we verified the interaction between parameters on a different dimensionality reduction technique. We observe the same effect when using UMAP (see Methods) instead of CMDS as a filter function (Supplementary **Fig. S7**).

In the case of real fMRI data, the resolution and gain parameters similarly affect the Mapper shape graph as we observe that three Mapper configurations have qualitatively different resulting shape graphs (**Fig. 7a-c**). When using a low gain value, the Mapper algorithm fails to extract the topological features of the input dataset because the shape graph does not connect the trajectory through the resting-state nodes (**Fig. 7a**). When the Mapper algorithm uses higher gain values, the shape graph has high connectivity patterns that lose the specificity of node types (**Fig. 7c**). In between those extreme values for the gain parameter, the Mapper algorithm shows the features expected: the resting-state nodes show a periphery trajectory while memory and math nodes are highly connected in the core hubs on the shape graph (**Fig. 7b**).

**Figure 7:**
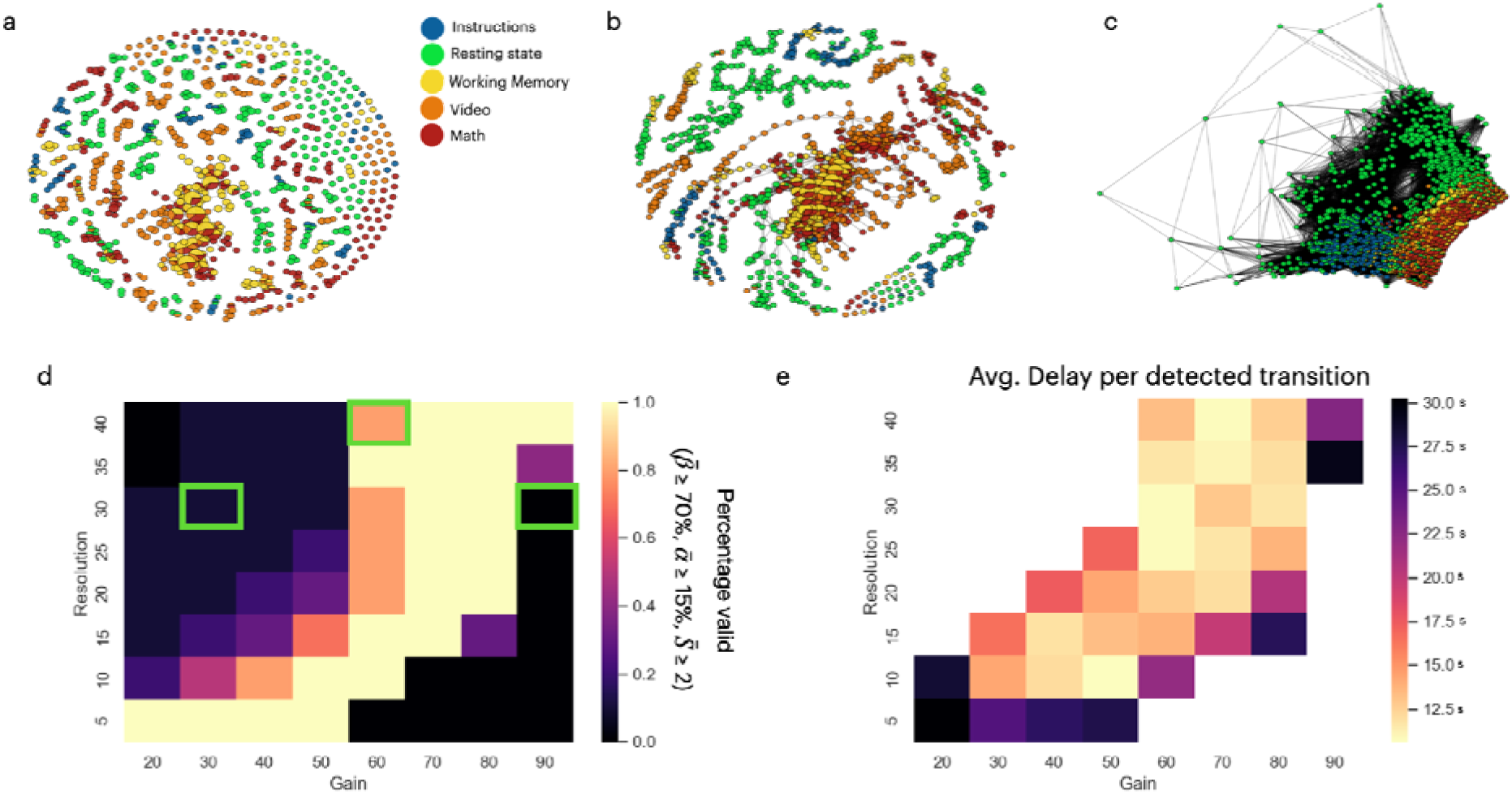
Choosing scale parameters when using the Mapper algorithm on fMRI data. **(a)** The Mapper shape graph on the fMRI dataset with an Extrinsic Mapper (resolution=30, gain=30%, k=12) for a single subject. Each node is represented by a pie chart of the composition of time points within that node. **(b)** An example of a Mapper shape graph with a discernible structure (resolution=40, gain=60%, k=12). **(c)** An example of an invalid shape graph result (resolution=30, gain=90%, k=12). All the Mappers shape graphs in this plot are generated with the geodesic Euclidean distance metric (needs the k parameter, k=12), CMDS embedding, extrinsic binning, and single linkage clustering. **(d)** Mapper configurations that pass the valid shape graphs criterion on a resolution-by-gain grid. The parameter varied is the k-value used to construct the geodesic distances. The green rectangles show the parameter configurations used for plots a, b, and c. **(e)** Heatmap of the delay per detected transition on a resolution-by-gain grid. The delay represents the average time (in seconds) between the transitions detected from the normalized degree and the ground truth of the actual transitions based on the event timings of each task. The missing values of the heatmap, represented by white colors (same as the background), are parameter configurations that return invalid Mapper shape graphs.

For a large number of parameter configurations (resolution, gain, and k parameters), the Mapper algorithm passes the criterion for a selection of appropriate resolution and gain parameters (**Fig. 7d**). As seen for the simulated dataset, the valid set of parameters are correlated for resolution and gain. For valid constructed graphs, the average delay of the predicted task transition spans from 3.2 to 39.4 seconds, with an average of 16.6 seconds (**Fig. 7e**). Accurate prediction of the transitions required a minimal value for resolution and gain (resolution > 5 and gain > 20%), which corresponds to the lower bound of the minimal number of nodes and connectivity required to represent the topology of the real dataset.

### 3.4 Effect of the clustering algorithm on the Mapper results

As the fourth step, the clustering method is essential for generating nodes in the final shape graph. We identify and analyze two clustering algorithms: Single Linkage and Density-based spatial clustering of applications with noise (DBSCAN) (**Fig. 8**). For the simulated dataset, the Linkage clustering method outperforms the DBSCAN algorithm by having, on average, more mapper shape graphs validated by the circleness GOF criterion (two-sample t-test: p < 0.05, **Fig. 8a**). Interestingly, for the real dataset, the DBSCAN algorithm outperformed the Single Linkage algorithm (two-sample t-test: p < 10^-5^, **Fig. 8b**). Breaking down the algorithms based on the hyperparameters used (**Fig. 8c**), we find a greater variation with the Single Linkage algorithm (ANOVA one-way, p < 10^-10^), while the DBSCAN algorithm has no variation in performance for its hyperparameters (ANOVA one-way, p > 0.05). Moreover, the best-performing hyperparameter configurations (single linkage bins=5, vs. DBSCAN eps=16) have no significant difference in performance (t-test, p > 0.05).

**Figure 8:**
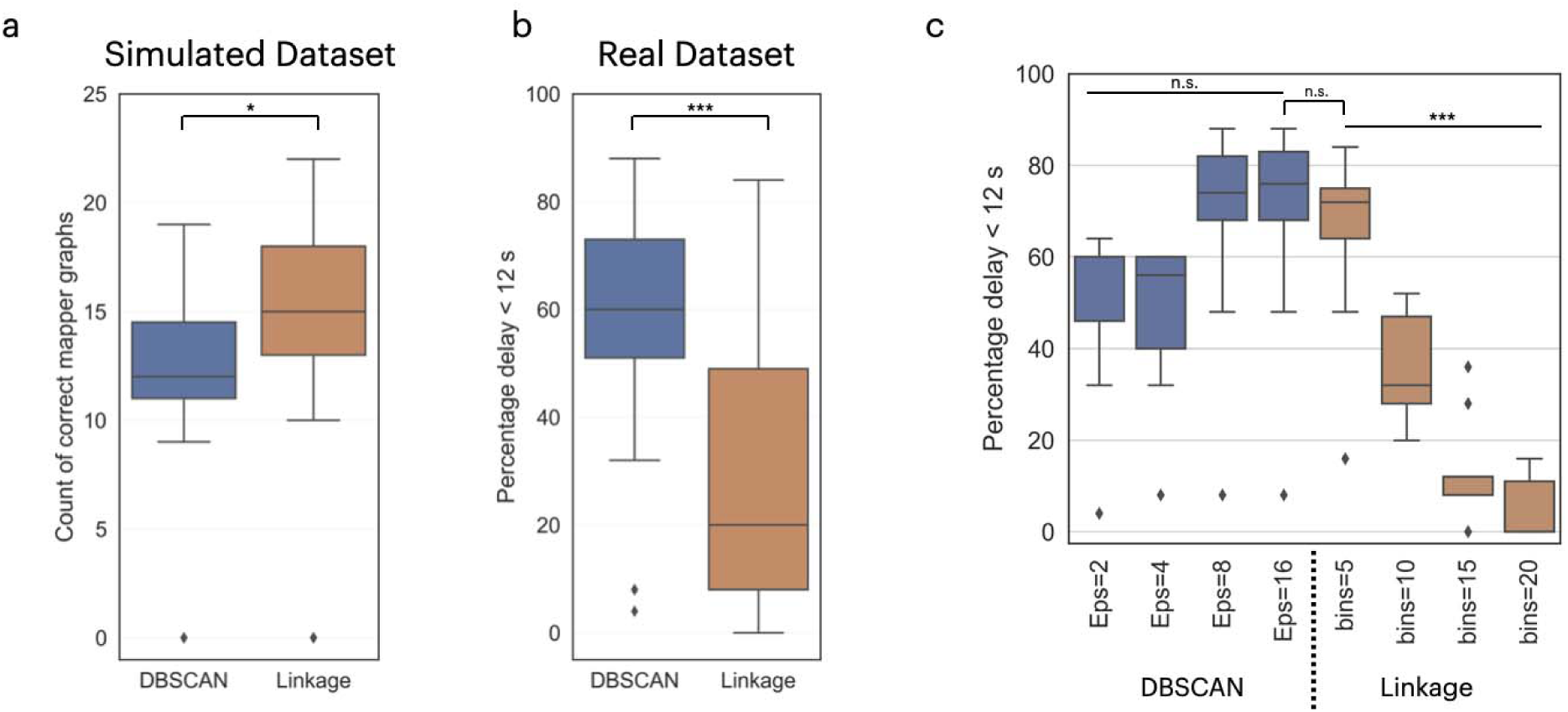
The effect of the clustering algorithm choices for the construction of the Mapper Shape graphs. **(a)** The performance of Mapper with different clustering methods on the simulated dataset. **(b)** The performance of Mapper using different clustering algorithms on the real dataset. **(c)** The breakdown of the performance of the clustering method is based on the hyperparameters chosen: the number of bins for the Single Linkage algorithm and the epsilon for the DBSCAN algorithm. The inverted-“U” connecting two distributions represents a t-test, and a straight line over multiple distributions represents an ANOVA test. The performance distributions in are shown as box plots. The result of the significance tests: n.s. is not significant p > 0.05; * is p < 0.05; ** is p < 0.01; *** is p < 0.001.

### 3.5 DeMapper software release

With the publication of this manuscript, we are releasing DeMapper as open-source software for use by the neuroscience community. We have developed the toolbox focusing on applications for neuroimaging datasets and use cases. Hence, the software can be used as either a MATLAB library that applies a single Mapper run or, alternatively, runs a batch of Mappers with different configurations (**Fig. 9**). The Mapper algorithm can be applied to any 2-dimensional matrix. In our neuroimaging applications, we have used Mapper on matrices representing measurements at different locations (parcels) over time, hence extracting the dynamical topology.

**Figure 9:**
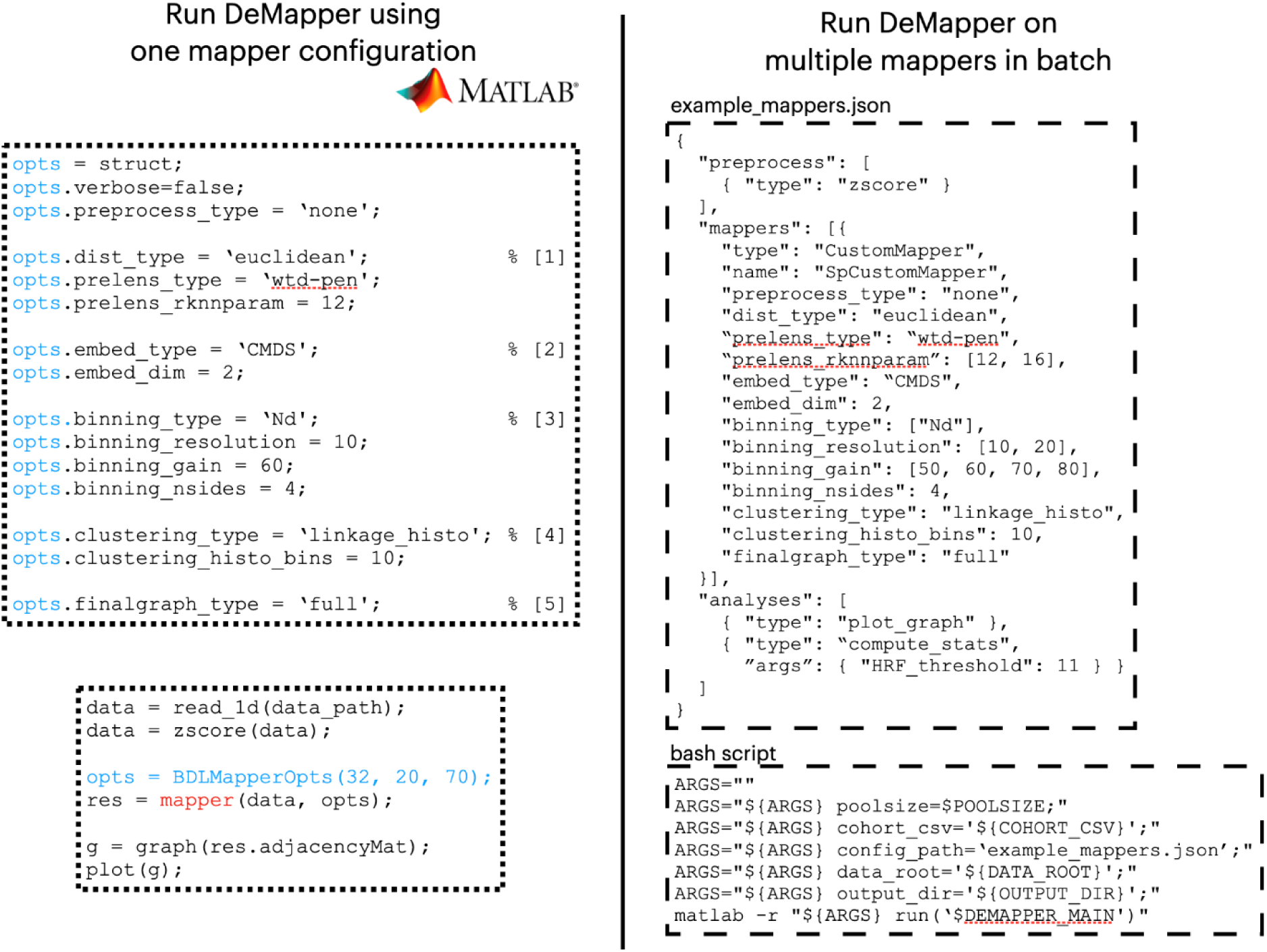
DeMapper code examples. **The left side** shows a code example for running DeMapper as one Mapper configuration and creating a simple graph out of it. **The top-left panel** shows how to set the parameters (opts) for a single configuration of Mapper: [1] picking the distance metric as a geodesic Euclidean distance metric with a k-value of 12; [2] picking the embedding algorithm as the CMDS embedding algorithm in 2 dimensions; [3] picking the binning strategy as the N-dimensional binning with 10 bins per dimension, a gain of 60%, and the bins are polygons with 4 sides (squares); [4] picking a clustering algorithm as the Linkage algorithm with 10 histogram bins; [5] generating a full graph with all the possible edges between nodes. **The bottom-left panel** runs the mapper configuration previously set on a dataset loaded and z-scored from ‘data_path.’ Moreover, it generates a simple graph based on the adjacency matrix of the mapper shape graph nodes. **The right side** shows code examples of running DeMapper on multiple parameter configurations. **The top-right panel** shows the configuration as written as a JSON file that describes the same parameters as in the mapper configuration set in the left side panel, with a few differences: the geodesic Euclidean distance will be tested with two k-values (12 and 16); the binning resolution will take two values (10 and 20 bins per dimension); the gain will take four values (50%, 60%, 70%, 80%); there are two extra analyses being run for each mapper generated (a plot graph and a compute stats with an HRF threshold of 11 seconds). This JSON configuration will generate a total of sixteen Mappers (two k-values by two resolutions by four gain parameters) with their associated analyses. **The bottom-right panel** shows how to run the DeMapper from a bash command to correctly reference the JSON configuration. The MATLAB variables defined are: poolsize determines the level of parallelization; cohort_csv is the path to a CSV file representing the inputs to be analyzed (subjects, sessions, etc.); config_path is the path to the JSON file describing the mapper configurations; data_root is the path to the input dataset, referenced relatively in the cohort_csv; output_dir is the path where to write the results. The sixteen mappers (defined by the JSON file) will be generated for each input to be analyzed (defined by the cohort CSV file).

An important feature of the DeMapper is customizing the Mapper parameters for fine-tuning based on each application. Furthermore, the batch analysis tool can scan multiple configurations to find the appropriate configuration for any input dataset, mimicking the hyperparameter search used in the Machine Learning field. Another useful feature for neuroimaging applications is providing a list of common presets for preprocessing and analysis. This feature requires a minimal setup for immediate plug-and-play usage on any dataset.

Moreover, with extra work, one can create custom extensions for preprocessing, analysis, and even Mapper steps, adapting the algorithm for each use case. As the number of Mappers scales with the number of configurations and inputs (subjects, sessions, etc.), the parallelization feature optimizes the runtime on multi-processor machines (**Fig. 9**). Finally, DeMapper was designed to be self-contained, requiring minimal installation steps so that the user can start using the package effortlessly.

## 4. Discussion

Despite the success of Mapper in uncovering brain dynamics during both resting and task-evoked states, there has been limited systematic investigation into the algorithm’s parameter selections and how they influence the resulting shape graphs. In this study, we analyzed various parameter choices for each deconstructed phase of the algorithm using synthetic and real fMRI datasets to comprehend their impact on the final shape graph depicting neural dynamics. Additionally, we briefly investigated the influence of noise on Mapper graphs and assessed their resilience when exposed to poor temporal resolution. As part of this research endeavor, we also released a Matlab-based toolbox, DeMapper, which could, in turn, facilitate convenient experimentation with Mapper, accommodating both naive and expert users. We hope this work could serve as a valuable resource for researchers (even beyond the field of neuroimaging) seeking to explore and analyze neural dynamics.

This paper provides several recommendations for researchers interested in utilizing the Mapper algorithm to analyze neuroimaging data, particularly fMRI data. First and foremost, based on our findings from simulated and real datasets, we prescribe using the Euclidean distance (ED) as the preferred distance metric. Previous studies have demonstrated the efficacy of ED in various applications involving neural data (Supekar, Musen, and Menon 2009; Kaiser 2011), such as fiber tracking in the brain (Kellmeyer and Vry 2016) and multivariate distance matrix regression analysis (Tomlinson et al. 2022). However, we acknowledge the need for future investigations to explore alternative distance measures. In particular, we hypothesize that angle measures, such as cosine similarity, may prove valuable in capturing higher-order interactions where geometric distances are unreliable.

Furthermore, our findings compel us to advocate using the geodesic distance metric construction based on the k-nearest neighbors (k-NN) algorithms. This approach to distance measurement captures the intrinsic local structure by encapsulating the correlation between subsequent steps of the time series. Given the propensity for neuroimaging datasets to exhibit pronounced interdependencies across successive temporal measurements, integrating geodesic metrics within the Mapper algorithm yields notable advantages in unraveling intricate patterns and dynamics inherent to such datasets.

In selecting a filter function for the Mapper algorithm, our investigation unveils insightful nuances when processing simulated and real neuroimaging datasets. Firstly, using Classical Multidimensional Scaling (CMDS) on constructed pairwise distances consistently reveals the correct topological shapes. Secondly, UMAP demonstrates an interesting effectiveness at low k-values, attributed to its own topological deconstruction process. Thirdly, despite preserving local structure, t-SNE struggles to capture the expected topological features. Drawing from these findings, we advocate adopting CMDS on geodesic pairwise distances as a robust choice for an extrinsic filter function within the Mapper algorithm. This configuration has proven successful across various applications in our endeavors (Saggar et al. 2022). Considering an alternative filter, the intrinsic Mapper (Geniesse et al. 2022), operating in the original space, showcases remarkable similarity in shape graphs to its extrinsic counterpart. The intrinsic approach even projects data into a space akin to high-dimensional embedding, enabling faster processing due to avoiding exponential bin proliferation. While the intrinsic Mapper represents a newer and more scalable version of the algorithm, our results suggest that the traditional extrinsic Mapper may be sufficient for analyzing simulated data and data derived from simple cognitive tasks, as employed in this study. However, we propose that future research explore the intrinsic Mapper’s potential advantages in analyzing complex task paradigms, such as naturalistic settings involving activities like watching movies or open-ended paradigms. Furthermore, considering the scalability of intrinsic Mapper, datasets other than neuroimaging, e.g., genetics which could contain millions of features and hundreds of thousands of rows, might be better suited for intrinsic Mapper.

Determining the optimal spatiotemporal scale for Mapper remains important in our research. The resolution and gain parameters are crucial in determining the level of detail in the resulting Mapper graphs, ranging from a single-node graph to having as many nodes as rows in the input data. Achieving scalability in representation has been a subject of extensive study, but there is no definitive answer yet. Thus, we recommend comprehensively exploring parameter choices across a broad range, potentially on a small sample of subjects (e.g., using a sandbox dataset for finetuning hyper-parameters). To enhance the search for optimal parameters, future studies could employ techniques like Bayesian hyperparameter tuning (Shahriari et al. 2016). Additionally, when reporting results, researchers should include parameter perturbation analyses to demonstrate the stability and reproducibility of their findings across various parameter choices. Moreover, an important future direction is investigating potential individual differences in Mapper binning parameters. It would be valuable to explore whether different subjects, age groups, or individuals with varying psychopathology profiles influence the spatiotemporal scale of brain dynamics, requiring further investigation and study.

Partial clustering is the fundamental step defining the Mapper algorithm, historically implemented through a single linkage (Singh et al. 2007). However, the rationale behind this preference instead of alternative methodologies lacks explicit justification. Our work underscores the need for further investigation to find the constraints for an optimal clustering algorithm, given that we revealed incongruent superior performers contingent on the dataset characteristics. While Single Linkage is conventionally favored in the context of Topological Data Analysis (TDA), we posit that a thorough evaluation of the DBSCAN algorithm is a potentially advantageous alternative.

Finally, some recommendations for reporting Mapper-generated results. First, validating the findings across multiple brain parcellations is advisable to ensure robustness and generalizability. This approach helps demonstrate that the observed patterns are consistent and not solely dependent on a specific parcellation scheme. Second, conducting parameter perturbation analyses is crucial for establishing the stability and reliability of the results across a wide range of parameter choices. This demonstrates that the findings are not mere artifacts of a particular parameter setting but reflect meaningful and consistent patterns in the data. Third, it is essential to employ appropriate null models, such as phase randomized null models, to account for linear and trivial properties of the data, such as autocorrelation in fMRI data. This allows for a more rigorous assessment of the significance of the observed patterns and helps distinguish genuine effects from random fluctuations. Finally, reporting individual-level results in addition to group averages is highly recommended. This individual-level analysis provides valuable insights into inter-subject variability and can reveal important nuances that might be obscured by averaging across participants.

### Limitations

Our study primarily focused on block design-based fMRI data, both simulated and real. However, it is essential to acknowledge that other fMRI experimental designs, such as event-related fMRI and naturalistic fMRI, present distinct challenges, and characteristics. The applicability and performance of the TDA-based Mapper approach in these alternative experimental designs remain uncertain. While recent research (Ellis et al. 2019) has shown promise in capturing topological structures from fast experimental designs, further investigation is warranted to evaluate the generalizability of our findings. Further, while we have presented empirical results illustrating the stability and reliability of Mapper graphs across a wide range of parameter configurations, we did not delve into the theoretical underpinnings of this stability. Prior studies have addressed the theoretical aspects of Mapper graphs (Bungula 2018, Carriere et al. 2018). Notably, recent work by Brown et al. (2021) explored the convergence of Mapper graphs in a probabilistic context. Future research should consider both empirical and theoretical aspects to provide a comprehensive understanding of Mapper graph stability. Our investigation is confined to fMRI data, and as such, our findings do not extend to other non-invasive human neuroimaging methodologies, such as EEG, fNIRS, and MEG. While Mapper has potential applications in invasive neuroimaging data, it may necessitate the exploration of different parameter configurations to accommodate the unique characteristics of these modalities. Future research should expand the scope to encompass a broader range of neuroimaging data sources. Lastly, an important limitation of our study lies in the comparison of different algorithms (e.g., UMAP, t-SNE) with varying parameter configurations. It is worth noting that each algorithm’s performance could potentially be improved with further fine-tuning. Our primary objective was to assess the ease of identifying suitable parameter configurations for accurate topological feature extraction. Future research can delve deeper into optimizing individual algorithms to refine their performance.

## Acknowledgments

This work was supported by an NIH Director’s New Innovator Award (DP2; MH119735), an NIH R01 MH127608, and an MCHRI Faculty Scholar Award to M.S.

