## Supplementary Information for "Deconstructing the Mapper algorithm to extract richer topological and temporal features from functional neuroimaging data"

a

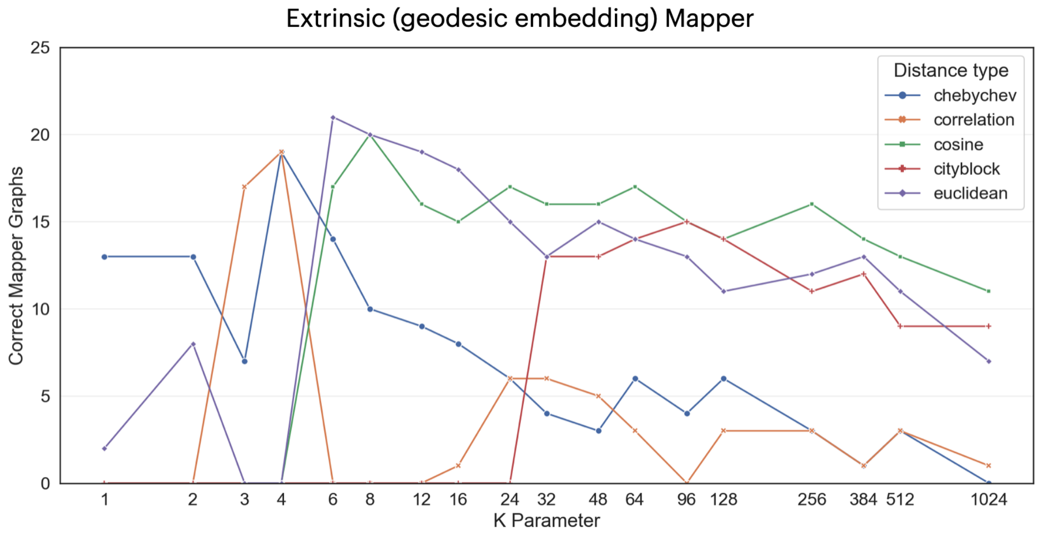


b

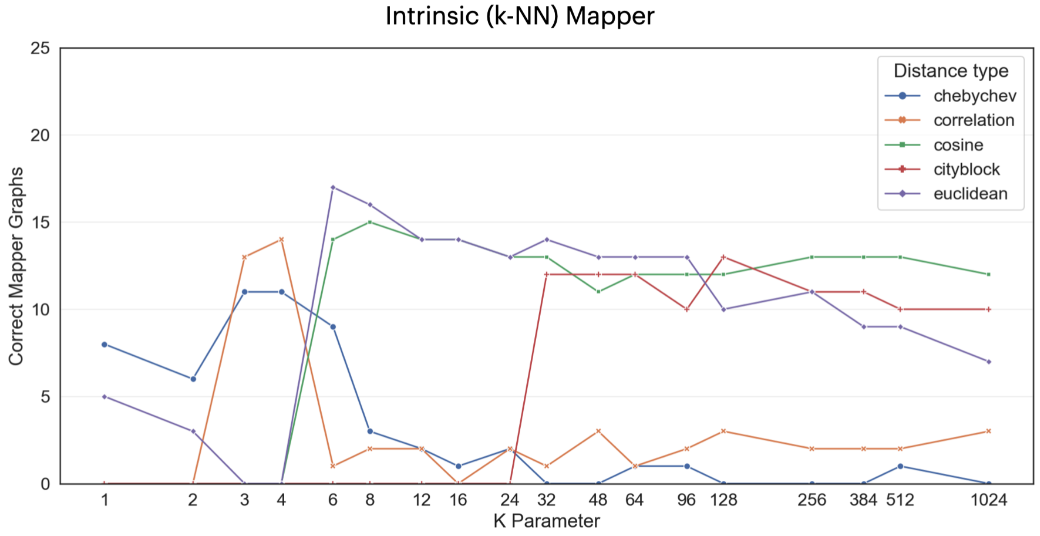


***Supplementary figure S1. The Mapper shape graphs performance on the simulated dataset, measuring based on the distance and k-value parameters.*** *The performance metric is the number of Correct Mapper shape graphs in a 5x5 resolution-by-gain grid (maximum 25) that capture the correct circle trajectory, as measured by the simulated dataset GOF measures (check Fig. 3a-b). We used different lenses:* ***(a)*** *an extrinsic Mapper (using geodesic distances and embeddings) and* ***(b)*** *an intrinsic mapper (working with the k-NN and avoiding dimensionality reduction).*

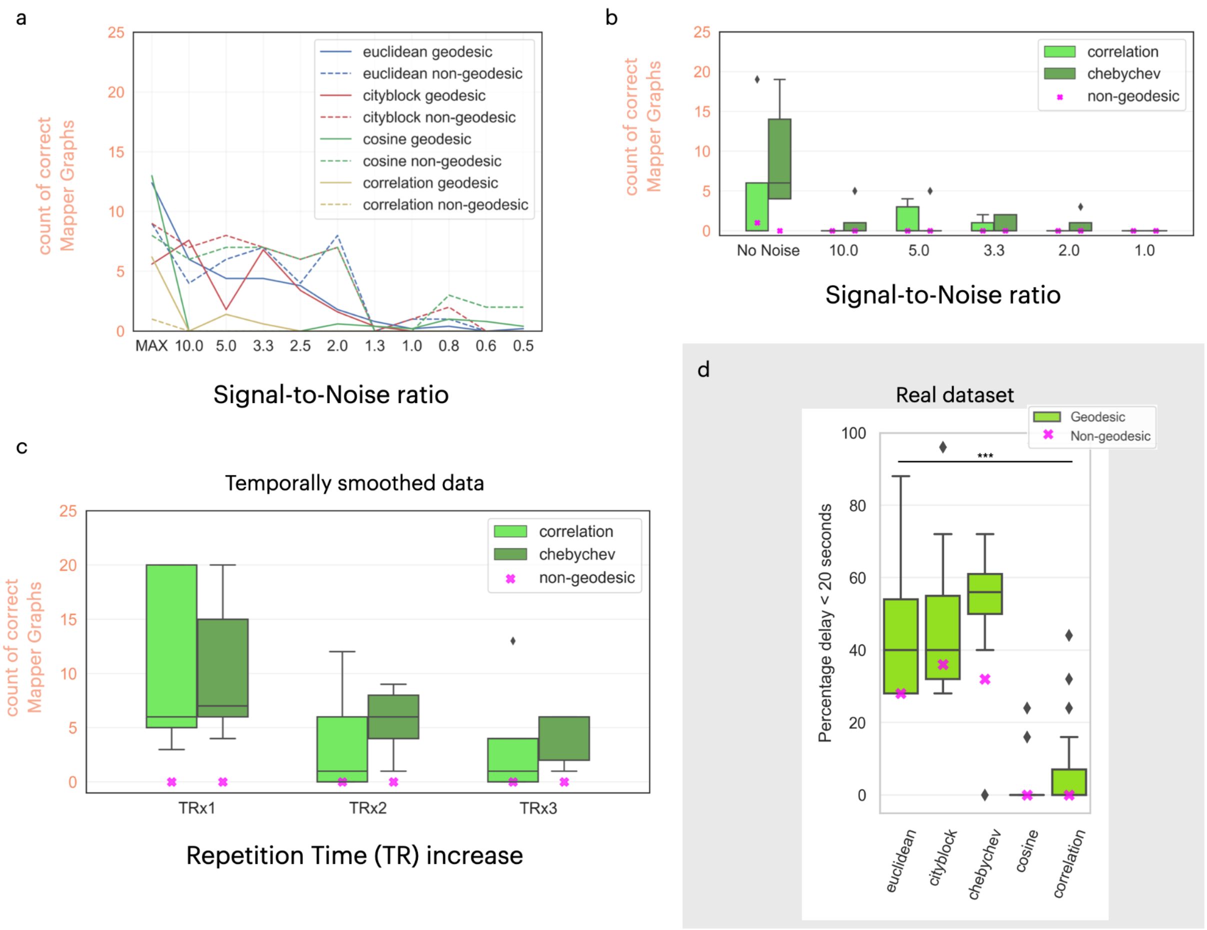


***Supplementary Figure S2. GOF metrics for different distance metrics for the simulated (a-c) and real (d) datasets. (a)** We show the performance of the Mapper algorithm on the simulated dataset as we decrease the signal-to-noise ratio (SNR). We plot the average performance for the 4 distance metrics (euclidean, cityblock, cosine, correlation) with and without geodesic distance metrics. **(b-c)** The equivalents plots of Fig. 3d and Fig. 3e with the remaining distance metrics (correlation and chebychev). **(d)** Replicating Fig. 4b with a higher delay threshold of 20 seconds. The line with *** denotes a one-way ANOVA with p < 10-10.*

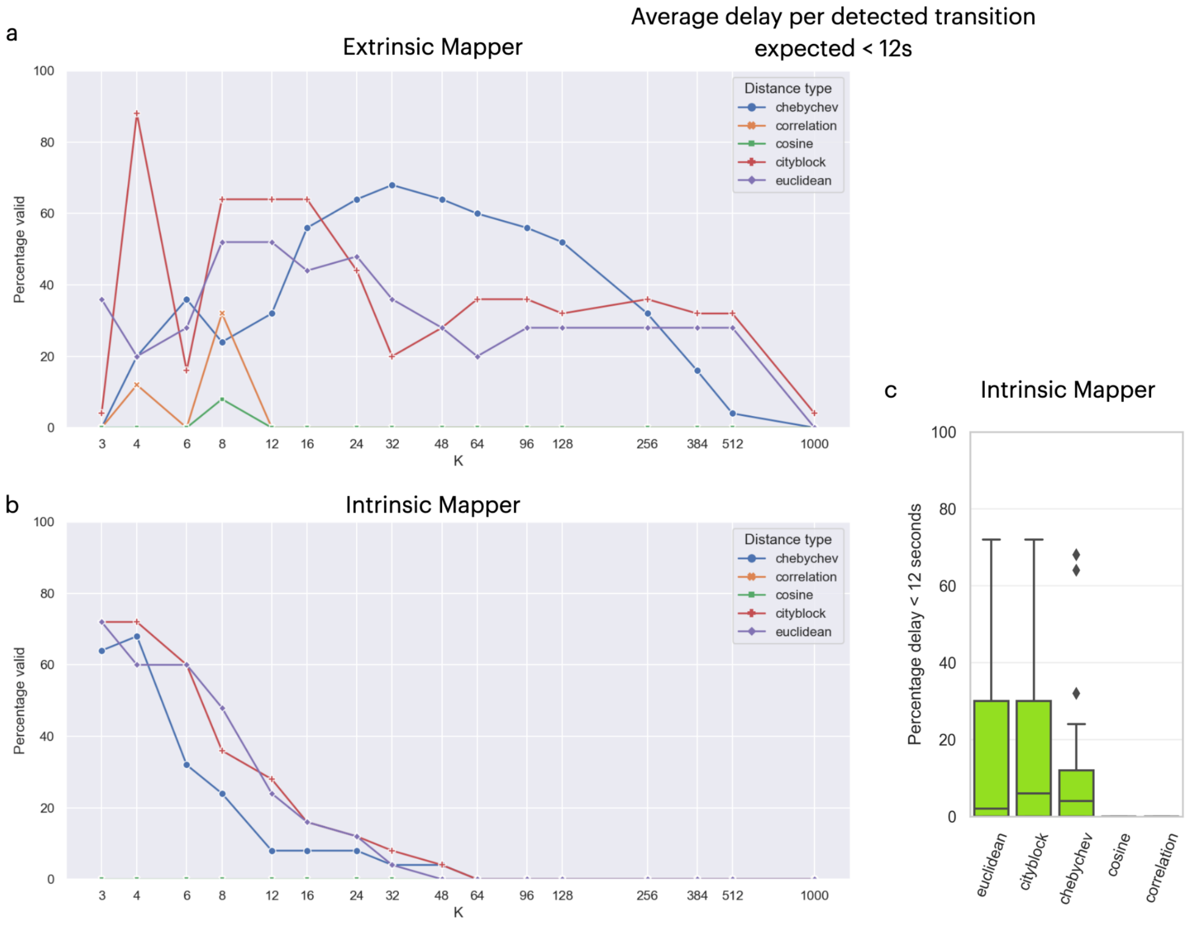


***Supplementary Figure S3. The Mapper shape graphs performance on the real dataset, measuring based on the distance and k-value used.*** *The performance metric is the percentage of valid Mapper shape graphs in a 5x5 resolution-by-gain grid (maximum 25) that pass the real dataset GOF criteria (check Methods and Fig. 4a). We used different lenses:* ***(a)*** *an extrinsic Mapper (using geodesic distances and embeddings) and* ***(b)*** *an intrinsic mapper (working with the k-NN and avoiding dimensionality reduction).* ***(c)*** *Aggregated results from plot (b). We used a 12-second delay threshold throughout this figure.*

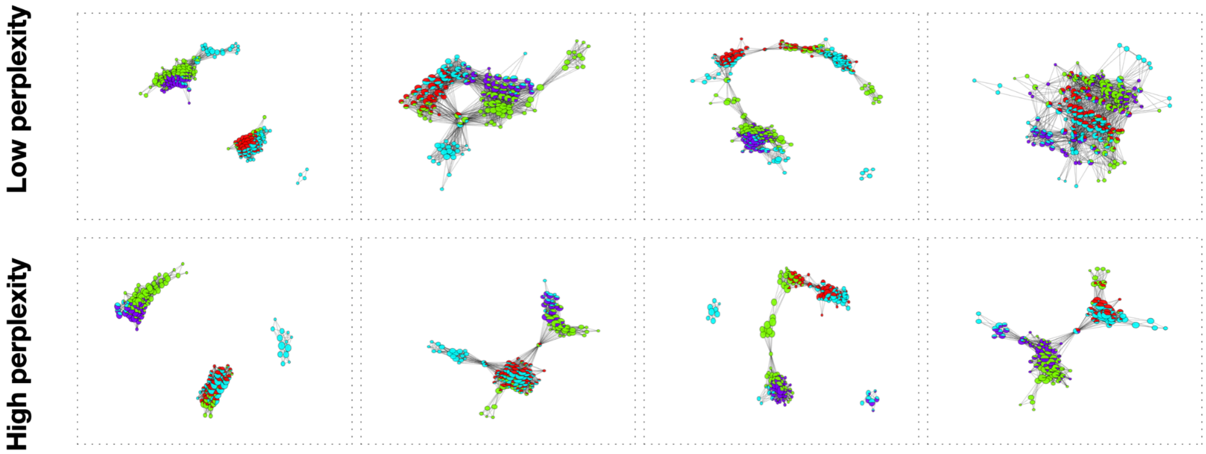


***Supplementary Figure S4. Example shape graph results using the t-SNE algorithm for dimensionality reduction, using different perplexity values.***

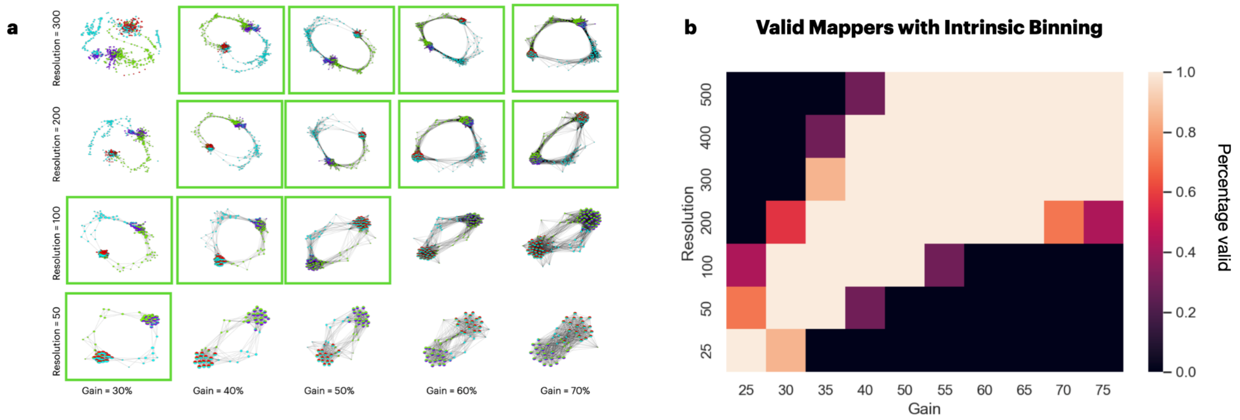


***Supplementary Figure S5. Performance of using different intrinsic binning parameters on the simulated dataset. (a)*** *Grid of shape graph produced by Intrinsic Mappers with k=20 for resolution parameters: 50, 100, 200, and 300; and gain parameters: 30%, 40%, 50%, 60%, and 70%. The valid Mapper results are highlighted within a green box.* ***(b)*** *A larger grid of valid Mapper results is aggregated over different k values. The plot shows three main regions, as seen in Fig. 6e, representing a band of resolution-gain parameter combinations that create valid Mapper graphs.*

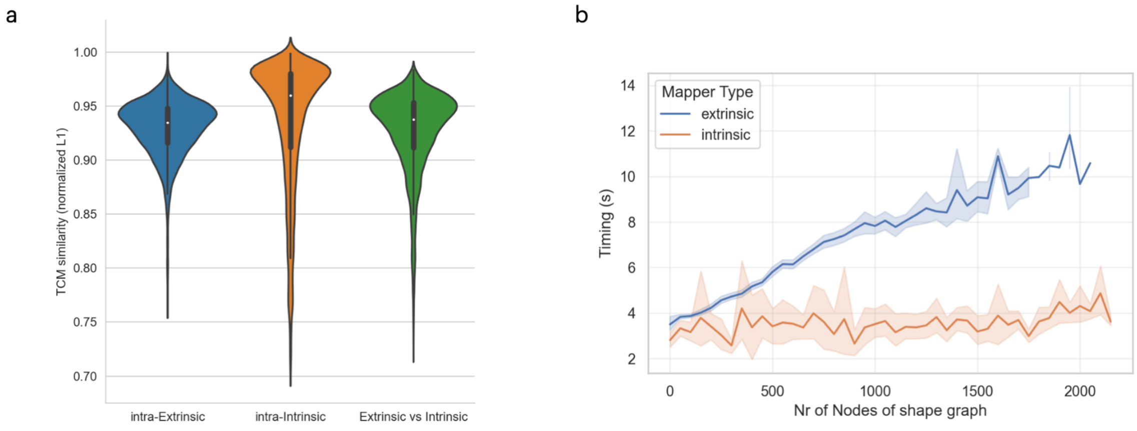


***Supplementary Figure S6. Comparison of Extrinsic and Intrinsic mapper graph representations and time performance.*** *(a) We computed the Temporal Connectivity Matrices (TCMs) generated by combinations of parameters for Extrinsic and Intrinsic Mappers. We computed the L1 similarity between all TCMs generated from the two types of mapper. Graphs produced by extrinsic and intrinsic Mappers show a high degree of similarity: 75% of comparisons are higher than 90% similarity (normalized L1) with a tail distribution spanning to a minimum of 70%. This distribution resembles the distribution of the extrinsic mapper's similarities. On average, intrinsic mappers have a higher similarity between the graphs generated.* ***(b)*** *For the extrinsic mapper's algorithm, the time it takes to compute a shape graph grows linearly with the number of nodes. The intrinsic mapper's algorithm has a constant performance as it handles the data as a graph instead of points in a grid of bins.*

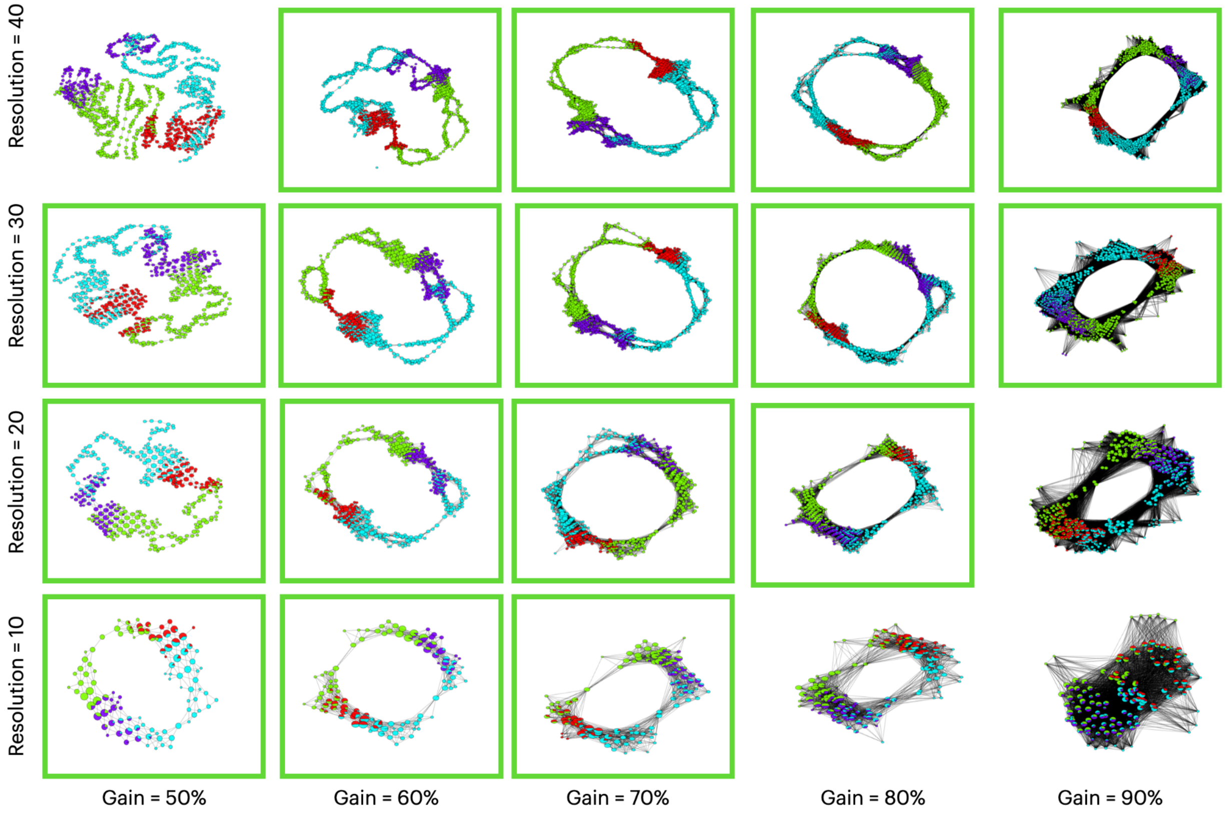


***Supplementary Figure S7. Example Mapper graphs generated using UMAP instead of CMDS for the Extrinsic Mapper algorithm.*** *The valid Mapper results are highlighted within a green box.*
